# The shape of density dependence and the relationship between population growth, intraspecific competition and equilibrium population density

**DOI:** 10.1101/485946

**Authors:** Emanuel A. Fronhofer, Lynn Govaert, Mary I. O’Connor, Sebastian J. Schreiber, Florian Altermatt

**Author notes:** **Correspondence Details**, Emanuel A. Fronhofer, Institut des Sciences de l’Evolution de Montpellier, UMR5554, Universit’e de Montpellier, CC065, Place E. Bataillon, 34095 Montpellier Cedex 5, France.

## Abstract

The logistic growth model is one of the most frequently used formalizations of density dependence affecting population growth, persistence and evolution. Ecological and evolutionary theory and applications to understand population change over time often include this model. However, the assumptions and limitations of this popular model are often not well appreciated.

Here, we briefly review past use of the logistic growth model and highlight limitations by deriving population growth models from underlying consumer-resource dynamics. We show that the logistic equation likely is not applicable to many biological systems. Rather, density-regulation functions are usually non-linear and may exhibit convex or both concave and convex curvatures depending on the biology of resources and consumers. In simple cases, the dynamics can be fully described by the continuous-time Beverton-Holt model. More complex consumer dynamics show similarities to a Maynard Smith-Slatkin model.

Importantly, we show how population-level parameters, such as intrinsic rates of increase and equilibrium population densities are not independent, as often assumed. Rather, they are functions of the same underlying parameters. The commonly assumed positive relationship between equilibrium population density and competitive ability is typically invalid. As a solution, we propose simple and general relationships between intrinsic rates of increase and equilibrium population densities that capture the essence of different consumer-resource systems.

Relating population level models to underlying mechanisms allows us to discuss applications to evolutionary outcomes and how these models depend on environmental conditions, like temperature via metabolic scaling. Finally, we use time-series from microbial food chains to fit population growth models and validate theoretical predictions.

Our results show that density-regulation functions need to be chosen carefully as their shapes will depend on the study system’s biology. Importantly, we provide a mechanistic understanding of relationships between model parameters, which has implications for theory and for formulating biologically sound and empirically testable predictions.

## Introduction

Population regulation and density dependence of population growth are at the core of fundamental but also controversial research in ecology (see e.g., Turchin, 1999; Henle et al., 2004; Sibly et al., 2005; Herrando-Prez et al., 2012; Krebs, 2015; Saether et al., 2016). Density dependence of population growth is often captured by the logistic growth model (Verhulst, 1838) and its more complex extensions, such as the θ-logistic model (Gilpin and Ayala, 1973). Despite its widespread use, it is important to recall that the logistic model is an abstract description of population dynamics (Herrando-Prez et al., 2012). This level of abstraction makes the interpretation of parameters challenging and may lead to paradoxical behaviours (Ginzburg, 1992; Gabriel et al., 2005; Mallet, 2012). Such issues are especially apparent in the often used *r – K* formulation with *r*_0_ being the intrinsic rate of increase, *K* the carrying capacity and *N* the population density:

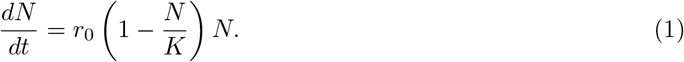

Additional challenges arise when these parameters, especially the carrying capacity (*K*), are interpreted in an evolutionary context (e.g., “*K*-selection” MacArthur, 1962). For instance, Luckinbill (1979) set out to test *r – K* selection theory using selection experiments in protist microcosms. Contrary to the expectation, he reported that r-selection actually led to higher carrying capacities, compared to the expected decrease in equilibrium population densities. Similar empirical evidence in the context of range expansions was reported by Fronhofer and Altermatt (2015) who showed that the interpretation of *K* as a parameter under selection and positively linked to competitive ability may be misleading. Recently, Reding-Roman et al. (2017) found positive *r – K* relationships in microbial systems counter to their initial hypothesis which led the authors to postulate ‘trade-ups’ and ‘uberbugs’ while discussing the relevance of these findings for cancer (see also Aktipis et al., 2013) and antibiotic resistance research (paper highlighted by Reznick and King, 2017). Although these and related issues have been discussed in detail by Matessi and Gatto (1984), Reznick et al. (2002), Rueffler et al. (2006) and Mallet (2012), to name but a few, current empirical work continues to expect negative *r – K* relationships (e.g., Fronhofer and Altermatt, 2015; Reding-Roman et al., 2017) and some theory continues to use “*K*” as an evolving trait (e.g., Lande et al., 2009; Burton et al., 2010; Fleischer et al., 2018, but see Engen and Sæther 2017).

In order to resolve some of the issues associated with the logistic growth model as described by Eq. 1, Mallet (2012), for instance, has promoted the use of Verhulst’s original r – *α* formulation of logistic growth (Verhulst, 1838, see also Kostitzin 1937). In comparison to the popular *r – K* formulation (Eq. 1), Verhulst’s model uses biologically interpretable parameters (Joshi et al., 2001; Ross, 2009; Mallet, 2012).

Namely, it includes *r*_0_ as the intrinsic rate of increase and *α* as the intraspecific competition coefficient:

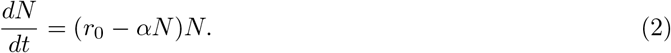

From Eq. 2 it follows that the population density at equilibrium is 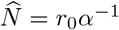. Similarly to Eq. 1, density dependence is assumed to act linearly, with *r*_0_ being the intercept (that is, the population growth rate when population density (*N*) is vanishingly small) and *α* representing the slope of population growth rate over population density.

Other authors have acknowledged the dynamic relationship between populations and their resources that causes density dependence by resorting to using more mechanistic consumer-resource models. For instance Matessi and Gatto (1984) show how resource dynamics and especially consumer traits have to be taken into account in order to understand density-dependent selection (see also Fronhofer and Altermatt, 2015). Such consumer-resource models provide a framework that can be used in an eco-evolutionary context (for a detailed discussion see McPeek, 2017) because model parameters linked to resource use (search efficiency, handling time) are related to real, individual-level traits that can be subject to evolutionary change (Rueffler et al., 2006; Govaert et al., 2019). Importantly, bottom-up population regulation due to renewing, depletable resources, as assumed in such consumer-resource models, is the most likely case according to Begon et al. (2006).

The disadvantage of these more mechanistic consumer-resource models is an increased complexity and number of parameters. Importantly, the quality and quantity of empirical data is often not sufficient for confronting such models to data. In an attempt to simplify, some studies have explored under what conditions population level growth models (e.g., the logistic, Eq. 1 or 2) can be used to describe the underlying consumer-resource dynamics. For instance, the consumer-resource dynamics underlying the logistic growth model have already been described by MacArthur (1970). A few years later, deriving the *r – K* logistic from the underlying resource dynamics, Schoener (1973) noticed that *r* and *K* share numerous parameters, implying that growth rates and resulting equilibrium densities may be linked through resource use traits (but see Getz, 1993). Similarly, Matessi and Gatto (1984) showed that selection for increased competitive ability (“K-selection”) does in fact not maximize equilibrium densities, but rather minimizes death rates and maximizes foraging rates and assimilation efficiencies. More recently, Abrams (2009b) compared the *θ*-logistic model (Gilpin and Ayala, 1973), an extension of the logistic model that allows for non-linear density dependence, to underlying dynamics in order to find plausible *θ* values (for an extension of this work to include multiple resources species see Abrams, 2009c). He showed that density dependence is non-linear if one assumes a Holling type II functional response for the consumer (see also Abrams, 2002), but that this non-linearity is different from the *θ* logistic model.

He finally extended his work to include a type III functional response and non-linear numeric responses. Reynolds and Brassil (2013) discussed, extended and reinterpreted these findings. Recently, O’Dwyer (2018) used the same approach to discuss the general applicability of Lotka-Volterra equations. Similar work deriving discrete-time population growth models has been conducted, for example, by Geritz and Kisdi (2004) and Brännström and Sumpter (2005). In parallel to this line of research on intra-specific density dependence, Lotka-Volterra models of inter-specific competition, and specifically the inter-specific competition coefficients (*α_i,j_*), have been linked to resource utilization through the concept of limiting similarity for example by MacArthur and Levins (1967), Schoener (1974) and Abrams (1975).

Here, we expand on this work and use consumer-resource models to derive different forms of population growth models and to gain a better understanding of their parameters, that is, how these parameters may be interpreted in biological terms and how they are inter-related. We focus on consumers in food chains that are bottom-up regulated, as described previously. Importantly, our considerations are mechanistic as we derive consumer density-dependent population growth without assuming that the resources grow logistically in the first place to avoid circularity in the argument (see the Supplementary Material S1 for a discussion of Lakin and Van Den Driessche, 1977). We furthermore investigate relationships between parameters that may constrain evolutionary trajectories. We use these derivations to show how intrinsic rates of increase, competitive abilities and equilibrium population densities are non-independent and provide explicit relationships between those parameters. In addition, we highlight the potential of deriving climate driven population growth models by discussing examples of how to extend our results to include temperature-dependence based on the metabolic theory of ecology (Brown et al., 2004) and multiple interacting species. Finally, we confront our theoretically derived growth models with time series data from microbial model populations using Bayesian inference.

## Modelling populations using consumer-resource models

In order to derive population growth models and the density-regulation function capturing how per capita consumer population growth rates (*r*) change depending on population density (*N*), as well as to understand relationships between population-level parameters, we will use the following general consumer-resource model in which *R* is the resource and *N* is the consumer population density:

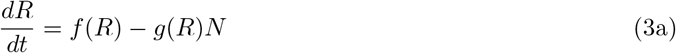

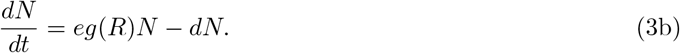

In this consumer-resource model, the function *f* (*R*) captures the growth of the resources and the function *g*(*R*) captures the functional response of the consumer (Holling, 1959), that is, how much resources are harvested by consumers depending on resource density. Furthermore, the constant *e* is the assimilation coefficient which translates consumed resources into consumer offspring and the constant *d* is the consumer’s death rate.

Using a time-scale separation argument (for a critical discussion see O’Dwyer, 2018), that is, assuming that resources quickly equilibrate 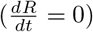 and solving Eq. 3a for *R*, we obtain the resource equilibrium density 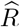 (piecewise defined, that is 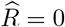 if there is no positive equilibrium). The per capita consumer dynamics (density-regulation function) then become:

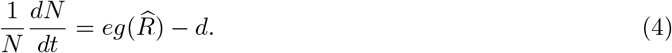

In order to understand how the form of the density-regulation function (Eq. 4) depends on different consumer and resource characteristics (e.g., filter feeders, saturating feeders, respectively, abiotic or biotic resources) we explore multiple realizations of *f*(*R*) and *g*(*R*), including a chemostat model for resource growth (*f*(*R*)) as well as linear and type II (saturating) functional responses for the consumer (*g*(*R*)). For an overview of the model components used below see Table 1. For simplicity, our work does not consider the possibility of predator-dependent functional responses (for a discussion and overview see Abrams, 2014), although these can be included in principle, for instance using a Beddington-DeAngelis functional response (Geritz and Gyllenberg, 2012). Such a functional response may also be able to represent spatial variation and behavioural complexities (Cosner et al., 1999).

**Table 1:**
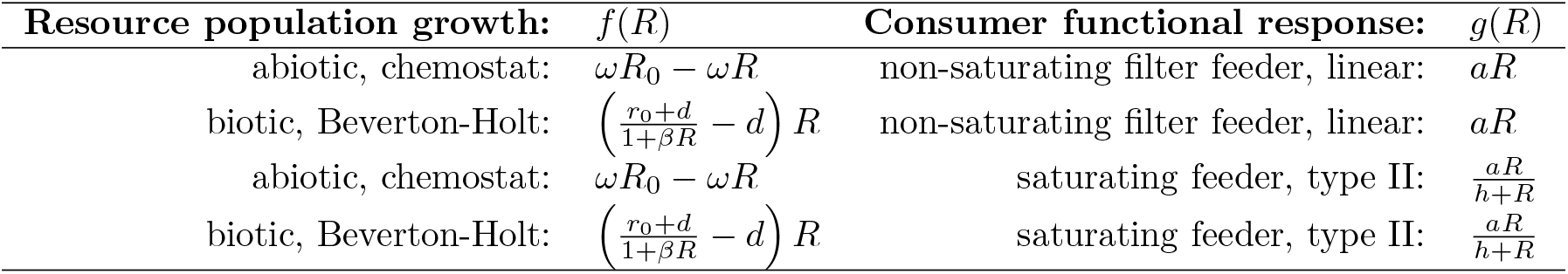
Model overview. Components and combinations of consumer-resource models used here. *R* is the resource population density. For chemostat resource population growth models *ω* is the flow rate into and out of the system and *R*_0_ is the resource concentration flowing into the system (we analyse an alternative chemostat formulation in the Supplementary Material S2). For biotic, that is Beverton-Holt, resource population growth models (for a justification of using this specific model, see main text), *r*_0_ is the intrinsic rate of increase, *d* the death rate and *β* the intraspecific competition coefficient. In consumer functional responses *a* represents the foraging rate an for saturating feeders, *h* is the half-saturation constant. Strictly speaking filter feeders will not exhibit purely linear functional responses, because also filter feeders have upper limits to filtering. We here use linear functional responses as approximations because of their mathematical tractability. If filter feeders are strongly limited in their resource intake capacity, our considerations using type II responses will be more appropriate.

| Resource population growth: | $f(R)$ | Consumer functional response: | $g(R)$ |
| --- | --- | --- | --- |
| abiotic, chemostat: | $\omega R_0 - \omega R$ | non-saturating filter feeder, linear: | $aR$ |
| biotic, Beverton-Holt: | $\left(\frac{r_0+d}{1+\beta R} - d\right) R$ | non-saturating filter feeder, linear: | $aR$ |
| abiotic, chemostat: | $\omega R_0 - \omega R$ | saturating feeder, type II: | $\frac{aR}{h+R}$ |
| biotic, Beverton-Holt: | $\left(\frac{r_0+d}{1+\beta R} - d\right) R$ | saturating feeder, type II: | $\frac{aR}{h+R}$ |

As a large part of the work introduced above, we stay in the realm of deterministic ordinary differential equations (ODEs), which have a long tradition in ecology and evolution. Of course, this implies that we are considering expected population sizes in continuous time systems and the equations may therefore be most appropriate for capturing the dynamics of biomass (Yodzis and Innes, 1992). Nevertheless, over finite time frames, trajectories of corresponding individual-based models are highly likely to remain close to trajectories of the ODE whenever habitat size is sufficiently large (Kurtz, 1981). Furthermore, ODEs can be derived as a deterministic approximation of stochastic processes describing the births and deaths of individuals, where interactions emerge by the “collision” of two individuals in a well-mixed environment (see also Mobilia et al., 2007).

We start by exploring density-regulation functions that are appropriate for basal biotic consumers feeding on abiotic resources. Then, we use these density-regulation functions to describe biotic resources fed upon by higher tropic level consumers and derive density-regulation functions describing the dynamics of the latter consumers. In all cases we show how population level parameters, specifically intrinsic rates of increase (*r*_0_) and equilibrium population densities 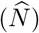, are impacted by and interrelated due to consumer traits such as parameters of the functional response (*g*(*R*)) as well as the assimilation coefficient (*e*) the consumer’s death rate (*d*).

## A simple case: non-saturating filter feeders

### Abiotic resources

For simplicity, we first assume that resources are abiotic, that is, resource population growth (*f*(*R*)) does not depend on resource population density, but rather on a fixed rate:

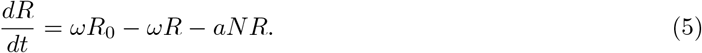

This resource model is often termed “chemostat model” with *ω* as the flow rate into and out of the system, and *R*_0_ as the resource concentration flowing into the system. The corresponding consumer dynamics are:

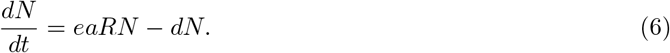

Chemostat models may follow different formulations and, for instance, assume no flow rate (see e.g., Abrams, 1988). This, however, does not impact our results qualitatively as the shape of the emerging density-regulation function as well as parameter relationships remain qualitatively unchanged (see Supplementary Material S2).

The amount of resources present at equilibrium can be obtained by setting Eq. 5 to zero, hence 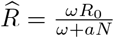. We can now substitute 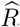 into Eq. 6 to obtain the per capita growth rate of the consumer:

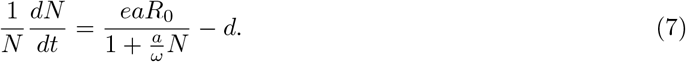

As Thieme (2003) notes, this result parallels the continuous-time version of the population growth model proposed by Beverton and Holt (1957) and derived by Schoener (1978) and Ruggieri and Schreiber (2005):

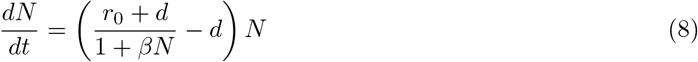

with *β* being the intraspecific competition coefficient in analogy to *α* in Verhulst’s model (Eq. 2). We will here always refer to the parameters scaling per capita population growth rate with population density as capturing intraspecific competitive ability.

In contrast to the logistic growth model (Eqs. 1 and 2) the density-regulation function described by Eq. 8 is not linear (see e.g., Pástor et al., 2016). Rather, it is convex, implying a decreasing strength of density regulation with increasing population density (Fig. 1; note that here and in the following a concave (convex) shape implies a (local) negative (positive) second derivative). Importantly, this implies that already for very simple consumer-resource systems the logistic growth model does not hold (but see Lakin and Van Den Driessche, 1977, and Supplementary Material S1 for a detailed discussion).

**Figure 1:**
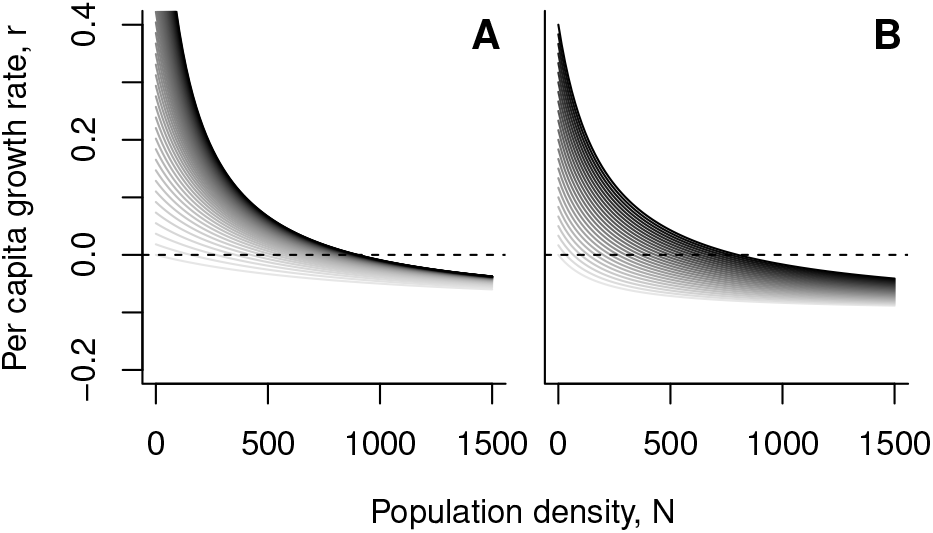
Density dependence for a non-saturating filter feeding consumer (chemostat model for the resource and linear functional response for the consumer; Eq. 7). (A) Effect of changing the foraging rate (*a*) while keeping the assimilation efficiency constant (*e* = 0.1). (B) Effect of changing the assimilation efficiency parameter (*e*) while keeping the foraging rate constant (*a* = 0.0005). In both panels, darker shades of grey indicate higher parameter values. Parameter examples: *R*_0_ = 10000, *ω* = 0.1, *e* ∈ [0.02, 0.1], *a*∈ [0.0001, 0.001], *d* = 0.1.

This derivation shows that non-saturating filter feeding consumer populations feeding on abiotic resources will generally have a convex per capita growth rate function (Eq. 7; Fig. 1) that is best described by a Beverton and Holt (1957) model (Eq. 8) and not by the logistic growth model given in Eq. 2. Following Jeschke et al. (2004), filter feeders that exhibit a type I functional response include branchiopods, some insect larvae, bryozoans, ascidians and molluscs, for example. Of course, the relevance of our derivation for any specific system depends on whether it fulfils relevant assumptions such that density is indeed regulated by food resources which may not be the case if densities are defined by spatial resources, for example. It is also important to note that we here assume a linear functional response for simplicity, and therefore no saturation of the consumer. Even if initially linear, functional responses of filter feeders will exhibit upper limits for high resource concentrations. Therefore, for clearly limited filter feeders, our considerations using type II functional responses below may be more appropriate.

We can also use Eq. 8 to study the relationship between population level parameters, such as the intrinsic rate of increase (*r*_0_) and the equilibrium density 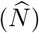 of the consumer population. For nonsaturating filter feeders the intrinsic rate of increase is

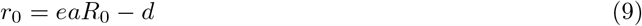

and the equilibrium density is obtained as

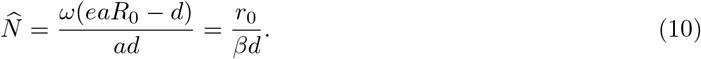

Importantly, this shows that intrinsic rates of increase (*r*_0_; Eq. 9), competitive abilities (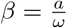; Eq. 8) and equilibrium population densities (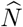; Eq. 10) depend on the same set of underlying parameters and are therefore not independent (see also Fig. 2). Competitive ability (*β*; Eq. 8) and intrinsic rates of increase (Eq. 9) are both linear functions of foraging rate (*a*). Note that the intrinsic rate of increase can nevertheless be independent of competitive ability, if the differences are driven by the assimilation efficiency (*e*). Counter to often made classical assumptions of a trade-off between growth rates and equilibrium densities, populations of consumers best characterized by this model will always exhibit a positive relationship between equilibrium densities 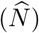 and intrinsic rates of increase, regardless of whether this is due to a change in foraging rate (*a*), assimilation efficiency (*e*) or death rate (*d*; see Fig. 2). The increase will be linear for *e*, concave for *a* and convex for *d*. If density-independent mortality (*d*) is very small, the effect of foraging rate (*a*) on the equilibrium population density 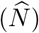 is negligible and 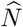 only depends on the assimilation efficiency (*e*).

**Figure 2:**
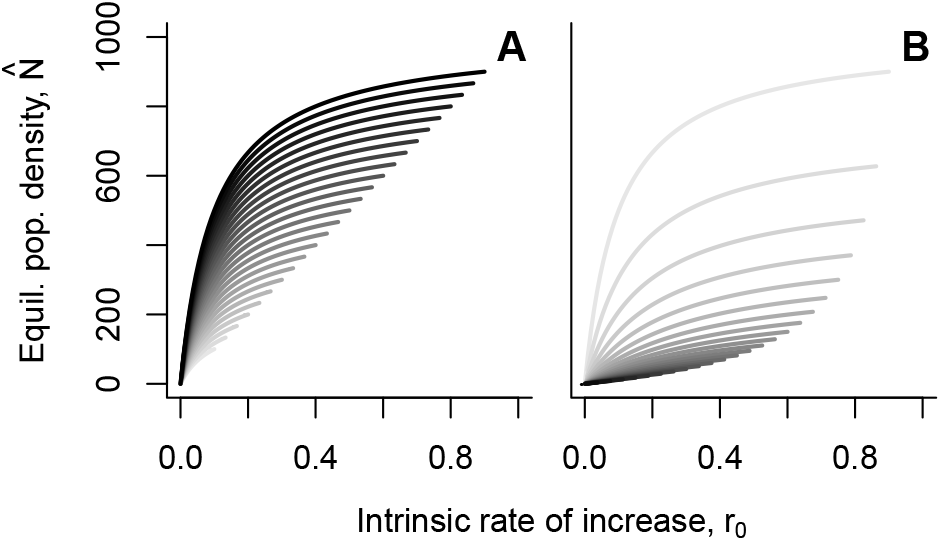
Relationship between intrinsic rate of increase (*r*_0_) and equilibrium population density 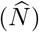 for consumers following a Beverton-Holt density-regulation function (chemostat model for the resource and linear functional response for the consumer; Eq. 7). (A) Effect of changing the foraging rate (*a*; solid lines; dots represent the *r*_0_ and 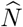 values for the highest value of *a*) and the assimilation efficiency (*e*; darker shades of grey indicate higher values of *e*) while the death rate is kept constant (*d* = 0.1). (B) Effect of changing the foraging rate (*a*; solid lines; dots represent the *r*_0_ and 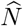 values for the highest value of *a*) and the death rate (*d*; darker shades of grey indicate higher values of *d*) while the assimilation efficiency is kept constant (*e* = 0.1). Parameter examples: *R*_0_ = 10000, *ω* = 0.1, *e* ∈ [0.02, 0.1], *a* ∈ [0.000 1, 0.00 1], *d* ∈ [0.1, 1].

### Biotic resources

Since the population dynamics of filter feeding consumers of a first trophic level can be described by the Beverton-Holt model (Eq. 8), we can investigate population dynamics of the next tropic level. We start by considering a non-saturating filter feeding resource where *f*(*R*) can be described by Eq. 8 and a non-saturating filter feeding consumer where *g*(*R*) is linear. Without repeating the previously described derivation, we can use the principle of the inheritance of the curvature described by Abrams (2009b), stating that the curvature of the density-regulation function of a consumer with a linear functional response is identical to the curvature of the density-regulation function of its resource. Hence, the population dynamics of any non-saturating filter feeding consumers in a food chain with a basal abiotic resource will follow the Beverton-Holt model (Eq. 8).

## A more complex case: saturating feeders

### Abiotic resources

Up to now we have assumed a linear functional response for *g*(*R*) which has been shown to be likely only applicable to some non-saturating filter feeding organisms (Jeschke et al., 2004). While keeping resources abiotic (*f*(*R*) as a chemostat model; Eq. 5), we next explore the form of the density-regulation function, as well as how intrinsic rates of increase, competitive abilities and equilibrium population densities are linked, if the consumer follows a saturating (type II) functional response (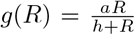, where *h* is the half-saturation constant).

Assuming resource equilibrium, we have to solve a quadratic equation (see Supplementary Material S3 for details). Substituting the resource equilibrium into the consumer equation results in a density-regulation function that is only slightly more complex than the Beverton-Holt model (Eq. 8; Fig. 1):

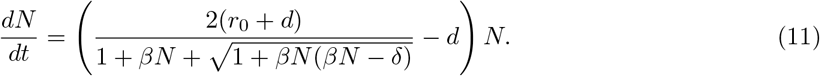

with 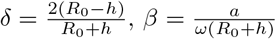 as the competitive ability, 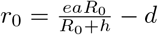 as the intrinsic rate of increase and

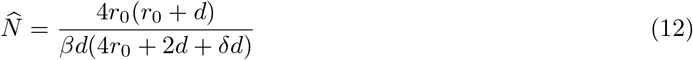

as the equilibrium density.

While the density-regulation function may now exhibit both concave and convex sections (Fig. 3, Eq. 11 and see Supplementary Material S4) it only depends on 4 parameters. The most relevant parameter driving the extent of the concave portion of the density-regulation function is the parameter *δ* and therefore the half-saturation constant (*h* in Eq. S14 and *δ* in Eq. 11; Fig. 3C). The density-regulation function is concave at low densities only if *δ* > 0 (which is equivalent to *R*_0_ > h; see Supplementary Material S4). The smaller the half-saturation constant, the more the concave part of the density-regulation function will approach a threshold-like shape. Ultimately the foraging rate becomes independent of consumer population density (*δ* → 2) which leads to exponential growth before density regulation kicks in at high densities (see Eq. S16). By contrast, the larger the half-saturation constant is, the more the type II functional response will approach a linear shape (*δ* → – 2). The latter case approaches the Beverton-Holt model (Eq. S17; see Supplementary Material S3 for details). As discussed in Supplementary Material S5, Eq. 11 behaves in part similarly to existing density-regulation functions, specifically the continuous-time version of the Maynard Smith and Slatkin (1973) model. However, these models do not have fewer parameters and therefore do not represent sensible alternatives.

**Figure 3:**
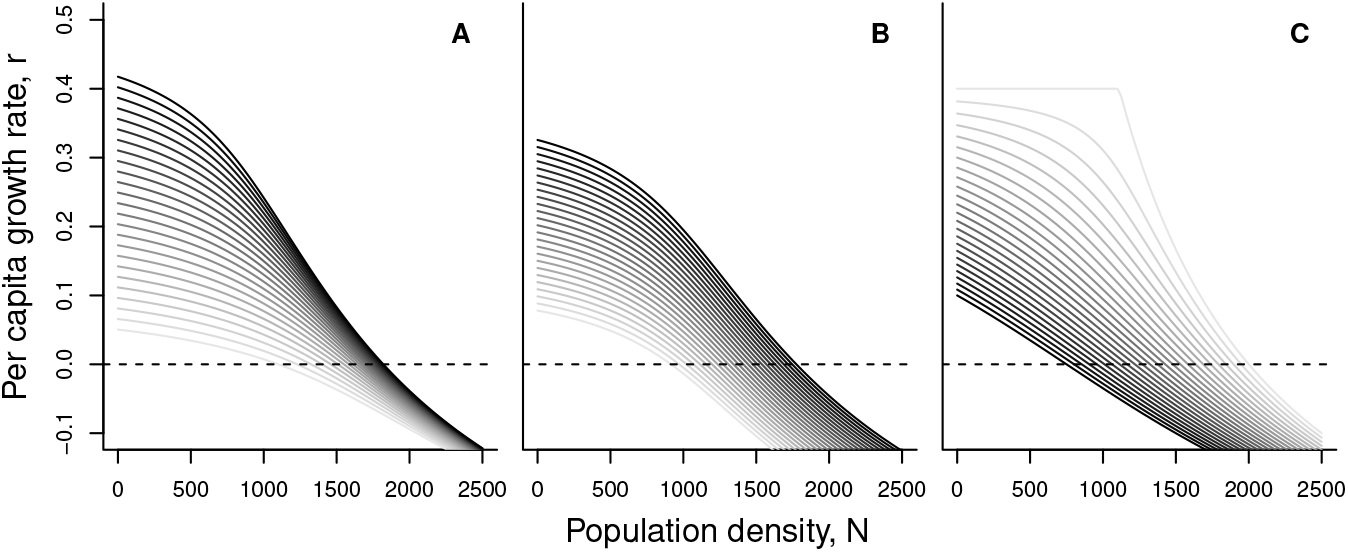
Density dependence for a consumer with a saturating, that is, type II, functional response (Eq. S14) while resource remain abiotic. (A) Effect of changing the foraging rate (*a*) while keeping the assimilation efficiency (*e* = 0.1) and the half-saturation constant (*h* = 9000) fixed. (B) Effect of changing the assimilation efficiency parameter (*e*) while keeping foraging rate (*a* = 9) and half-saturation constant (*h* = 9000) fixed. (C) Effect of changing the half-saturation constant (*h*) while keeping foraging rate (*a* = 9) and assimilation efficiency (*e* = 0.1) fixed. In all panels darker shades of grey indicate higher parameter values. Parameter examples: *R*_0_ = 100000, *ω* = 0.1, *e* ∈ [0.07,0.1], *a* ∈ [6,10], *h* ∈ [0, 50000], *d* = 0.5.

As in the simplest case of non-saturating filter feeders, intrinsic rates of increase (*r*_0_; Eq. S12), competitive abilities (*β*; Eq. S15) and equilibrium population densities (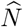; Eq. 12) depend on the same set of underlying parameters and are therefore not independent. All three parameters increase with the maximal foraging rate (*a*) and decrease with the half-saturation constant (*h*) and death rate (*d*). While competitive ability does not depend on the assimilation efficiency (*e*), equilibrium population densities and population growth rates increase with increasing assimilation efficiency. Therefore, as for non-saturating filter feeders, relationships between equilibrium population density 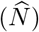 and intrinsic rates of increase (*r*_0_) will always be positive although the exact shape of the function will depend on which of the underlying consumer parameters is changing (Fig. 4).

**Figure 4:**
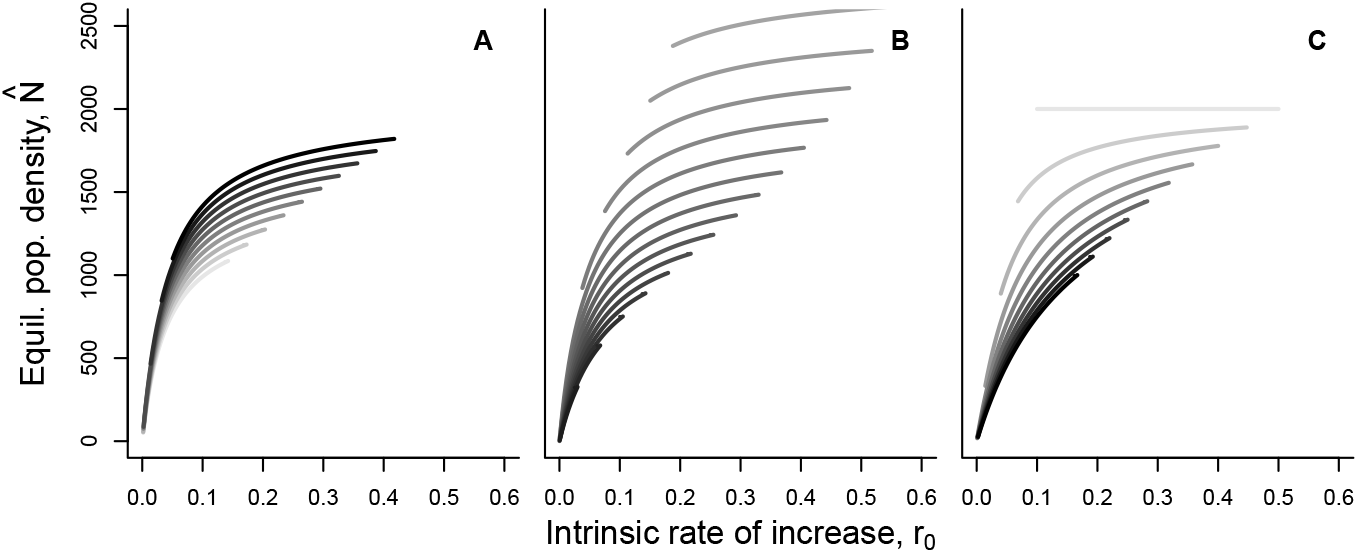
Relationship between intrinsic rate of increase (*r*_0_) and equilibrium population density 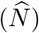 for consumers following Eq. 11 (chemostat model for the resource and saturating, that is, type II functional response for the consumer). (A) Effect of changing the foraging rate (*a*; solid lines; dots represent the *r*_0_ and 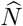 values for the highest value of *a*) and the assimilation efficiency (*e*; darker shades of grey indicate higher values of *e*) while the half-saturation constant and the death rate are kept constant (*h* = 9000, *d* = 0.5). (B) Effect of changing the foraging rate (*a*; solid lines; dots represent the *r*_0_ and 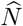 values for the highest value of *a*) and the death rate (*d*; darker shades of grey indicate higher values of *d*) while the half-saturation constant and the assimilation efficiency are kept constant (*h* = 9000, *e* = 0.1). (C) Effect of changing the foraging rate (*a*; solid lines; dots represent the *r*_0_ and 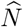 values for the highest value of *a*) and the half saturation constant (*h*; darker shades of grey indicate higher values of *h*) while the death rate and the assimilation efficiency are kept constant (*d* = 0.5, *e* = 0.1). Parameter examples: *R*_0_ = 100000, *ω* = 0.1, *e* ∈ [0.07,0.1], *a* ∈[6,10], *h* ∈ [1,50000], *d* ∈ [0.1,1].

### Biotic resources

Consumers can also exhibit saturating functional responses (*g*(*R*)), while feeding on biotic resources. In this case Eqs. S5a and S5b have to be adjusted as shown in the Supplementary Material S6 (Eqs. S27a–S27b) which yields an even more complex density-regulation function (Fig. 5).

**Figure 5:**
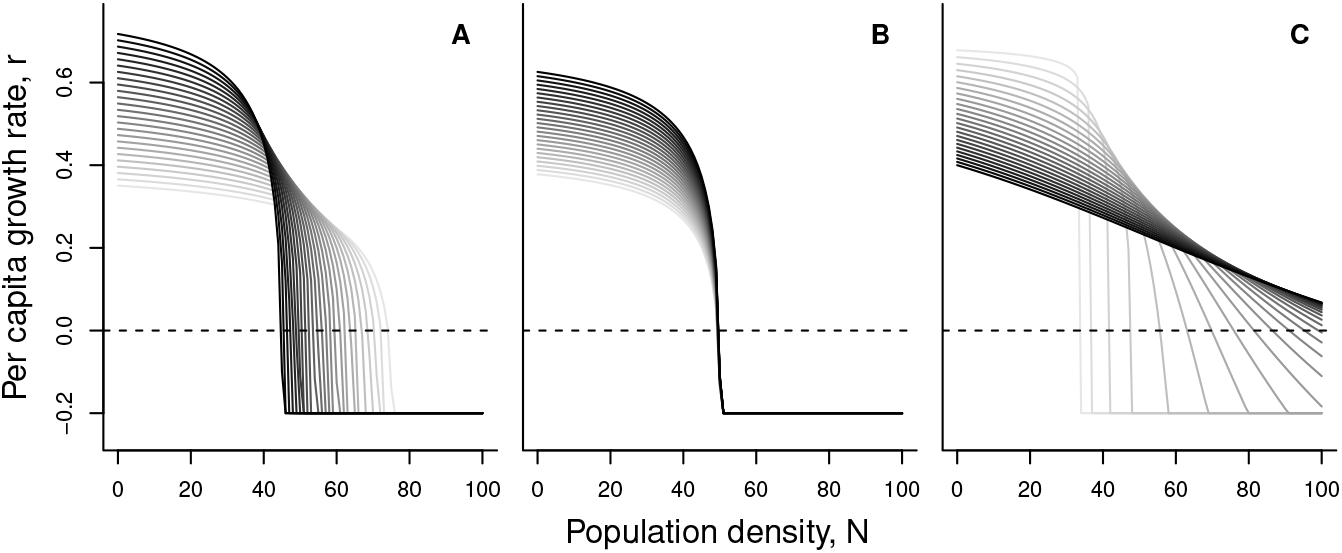
Density dependence for a consumer in a consumer-resource system with biotic resources (following the Beverton-Holt model; Eq. 8) and a saturating, that is, type II, functional response. (A) Effect of changing the foraging rate (*a*) while keeping assimilation efficiency (*e* = 0.1) and the half-saturation constant (*h* = 900) fixed. (B) Effect of changing the assimilation efficiency parameter (*e*) while keeping foraging rate (*a* = 9) and half-saturation constant (*h* = 900) fixed. (C) Effect of changing the half-saturation constant (*h*) while keeping foraging rate (*a* = 9) and assimilation efficiency (*e* = 0.1) fixed. In all three panels, darker shades of grey indicate higher parameter values. Parameter examples: *r*_0,*R*_ = 0.5, *β_R_* = 0.001, *d_R_* = 0.05, *e* ∈ [0.07,0.1], *a* ∈[6,10], *h* ∈ [250,5000], d= 0.2.

While some simplifications are possible, the resulting density-regulation function remains unwieldy with its 7 parameters (see Eq. S37). Clearly, if such complex dynamics need to be analysed in detail, it might be more appropriate to directly rely on the consumer-resource model. However, Eq. S37 is a density-regulation function that, interestingly, behaves similarly to the well known, continuous-time version of the Maynard Smith and Slatkin (1973) model (see Supplementary Material S7 and Fig. S5):

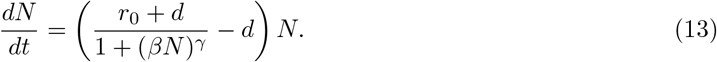

In this model, the equilibrium density is 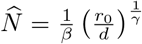. Via its shape exponent *γ* (Eq. 13) the Maynard Smith-Slatkin model is flexible enough (Bellows, 1981) to reproduce both the convex and concave parts of the density-regulation function (see Fig. S5 and S6) qualitatively. It is however not able to capture aspects like the asymmetry well (see Fig. S5).

Up to here, for abiotic and biotic resources and non-saturating filter feeding consumers, the equilibrium density was always a monotonically increasing function of the intrinsic rate of increase. In contrast to these simpler consumer-resource systems, our analyses show that for saturating consumers feeding on biotic resources this relationship can take many forms: unimodal, monotonically decreasing or increasing and 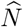 and *r*_0_ can even be independent of each other (Fig. 6). Changes in the foraging rate (*a*) may lead to positive or negative relationships which are globally unimodal and concave (Fig. 6 A). Interestingly, changes in the assimilation efficiency (*e*) may not impact 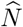 while increasing *r*_0_ (Fig. 6 A). Decreasing death rates (*d*) lead to a monotonically increasing relationship between 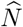 and *r*_0_ (Fig. 6 B). Clearly, the unimodal pattern is centrally impacted by the half-saturation constant (*h*; Fig. 6 C): very low values of *h* can lead to negative relationships between 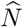 and *r*_0_. Such a negative relationship is otherwise only possible in the Lakin and Van Den Driessche (1977) model (see Supplementary Material S1).

**Figure 6:**
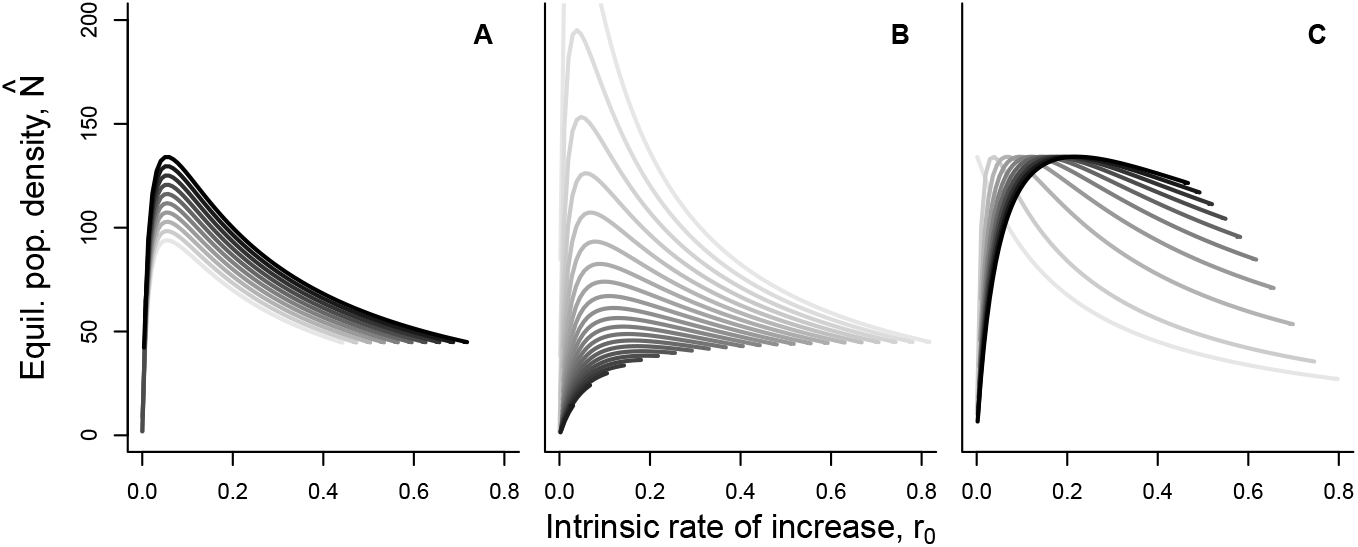
Relationship between intrinsic rate of increase (*r*_0_) and equilibrium population density 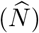 for saturating consumers feeing on biotic resources, that is, following Eq. S37 (Beverton-Holt model for the resource and saturating, that is, type II functional response for the consumer). (A) Effect of changing the foraging rate (*a*; solid lines; dots represent the r_0_ and 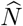 values for the highest value of *a*) and the assimilation efficiency (*e*; darker shades of grey indicate higher values of *e*) while the half-saturation constant and the death rate are kept constant (*h* = 900, *d* = 0.2). (B) Effect of changing the foraging rate (*a*; solid lines; dots represent the *r*_0_ and 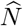 values for the highest value of *a*) and the death rate (*d*; darker shades of grey indicate higher values of *d*) while the half-saturation constant and the assimilation efficiency are kept constant (*h* = 900, *e* = 0.1). (C) Effect of changing the foraging rate (*a*; solid lines; dots represent the *r*_0_ and 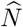 values for the highest value of *a*) and the half saturation constant (*h*; darker shades of grey indicate higher values of *h*) while the death rate and the assimilation efficiency are kept constant (*d* = 0.2, *e* = 0.1). Parameter examples: *r*_0,*R*_ = 0.5, *β_R_* = 0.001, *d_R_* = 0.05, *e* ∈ [0.07, 0.1], *a* ∈ [0,10], *h* ∈ [25,5000], *d*∈ [0.1,1].

## Confronting population growth models with data

Above, we show that many consumers that are bottom-up regulated will exhibit non-linear density regulation (Figs. 1, 3 and 5). For capturing dynamics at the consumer level, we offer alternatives to the logistic growth formulation: In the simplest case of non-saturating filter feeders and abiotic resources the dynamics of the consumer follow exactly the Beverton-Holt model (Eq. 8). For more complex cases involving saturating feeders (non-linear functional responses) or biotic resources Eq. 11 holds or the Maynard Smith-Slatkin model (Eq. 13) can be used as a heuristic description.

We will now consider how well these models perform when confronted with empirical data of a real biological system. We used populations of the freshwater ciliate model organism *Tetrahymena thermophila* as a consumer that feeds on the bacterium *Serratia marcescens*. Starting at low population densities, we recorded population growth trajectories over the course of two weeks for 7 different genotypes, each replicated 6 times. We fit the logistic, the Beverton-Holt model, Eq. 11, and the Maynard Smith-Slatkin model to these dynamics using a Bayesian approach (trajectory matching; i.e., assuming pure observation error; see Supplementary Material S8 and Rosenbaum et al. (2019) for details) to avoid the pitfalls of likelihood ridges (Clark et al., 2010; Delean et al., 2012) and compared fits using WAIC as an information criterion (McElreath, 2016).

Given our theoretical considerations, as well as the fact that the bacterial resources were replenished regularly in the microcosms (effectively mimicking abiotic resource flows), we expect either the Beverton-Holt model (BH) or Eq. 11 to best describe the dynamics and therefore produce the best fit to the time-series data compared to other candidate models of density-dependent dynamics. Indeed, over the 42 growth curves the logistic model had, on average, the worst fit (mean WAIC over all fits = 5.68; for individual results see Fig. S7 and Tab. S1), followed by the Beverton-Holt model (mean WAIC = 0.64) and the Maynard Smith-Slatkin model (mean WAIC = 0.22) while Eq. 11 performed best on average (mean WAIC = −0.16; see Fig. 7 for an example and Fig. S7 for all fits). In only 3 out of the 42 time-series the logistic was found to fit best (Fig. S7). The remaining growth curves were found to follow most often Eq. 11 (26 out of 42), the Beverton-Holt model (7 out of 42) or the Maynard Smith-Slatkin model (6 out of 42). This finding is in good accordance with work showing that *Tetrahymena may* follow a type II functional response (Fronhofer and Altermatt, 2015).

**Figure 7:**
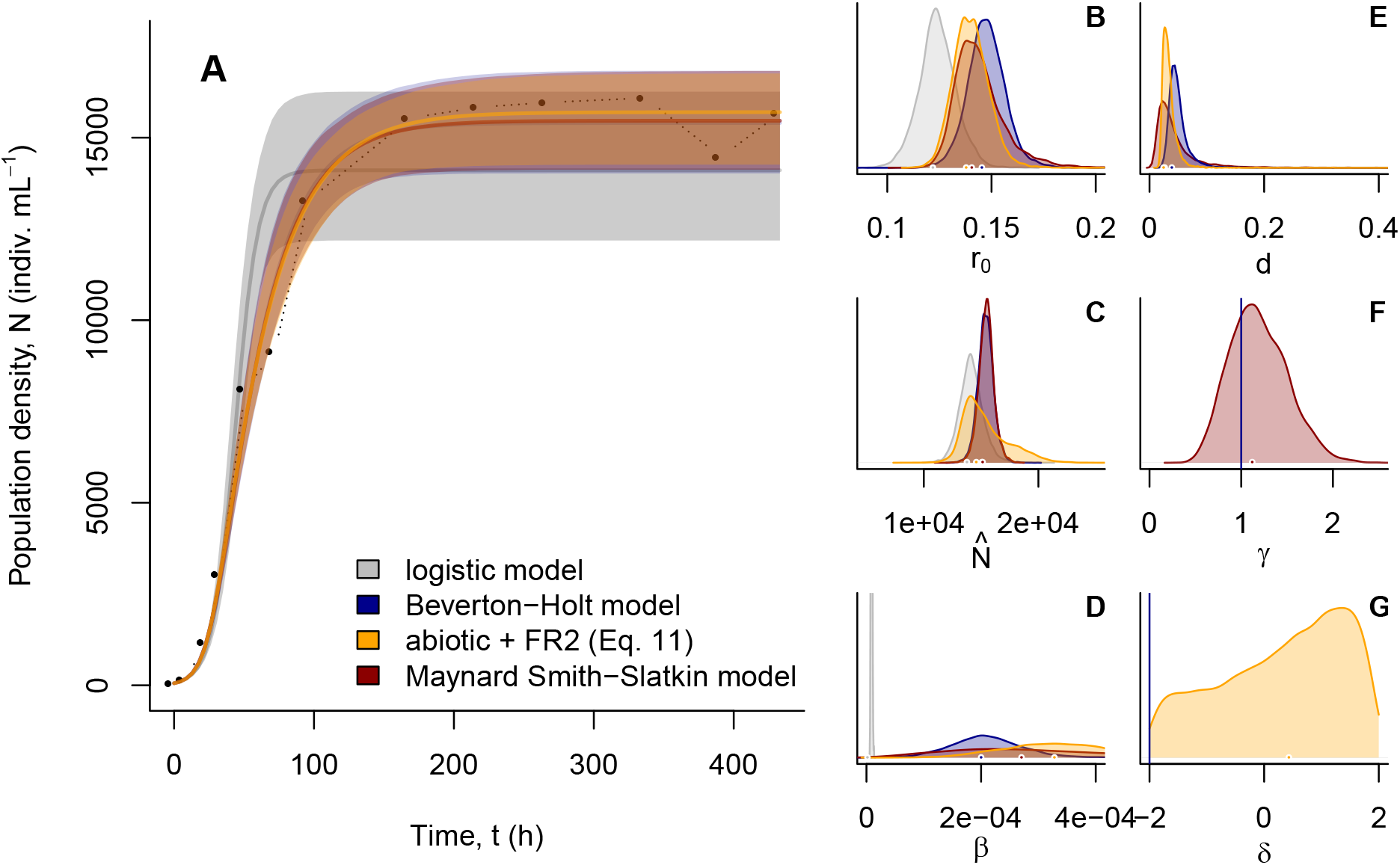
Fitting population growth models to *Tetrahymena thermophila* dynamics using Bayesian inference (see Supplementary Material S8 and Rosenbaum et al. (2019) for details). We fitted the logistic (black; Eq. 1), the Beverton-Holt model (blue; Eq. 8), Eq. 11 (orange), as well as the Maynard SmithSlatkin model (red; Eq. 13) and compared fits using WAIC. Eq. 11 fitted best (WAIC = −19.21), but was very similar to both the Maynard Smith-Slatkin model (WAIC = −18.12) and to the Beverton-Holt model (WAIC = −16.81) while the logistic model clearly performed worst (WAIC = −2.38). As becomes clear in panel (A) the logistic model was not able to capture the asymmetry of the empirical growth curve well that approaches the equilibrium density more slowly than the logistic allows for (see also Fig. 8 A). We report medians and the 95th percentile of the posterior predictive distribution. Panels (B-G) show the posterior distributions of the parameters of the four models. Because resource dynamics were very strictly controlled, we predicted that this system follows abiotic resource dynamics, which is in good accordance with the fit of Eq. 11 and the low estimate of *γ* (see F and Fig. S4). As a consequence, *Tetrahymena thermophila* likely exhibits a type II functional response which is in good accordance with work by Fronhofer and Altermatt (2015).

## Implications and extensions

Our derivations and the empirical support for Eq. 11 and the Beverton-Holt model clearly show the value of considering alternatives to the logistic growth model when modelling population dynamics with density-dependent rates. Even more importantly, our work highlights the underlying consumer-resource parameters being responsible for changes of and relationships between population level growth parameters (Figs. 2, 4 and 6). In the following section we discuss evolutionary consequences of our findings, and showcase extensions of our models to include temperature-dependence or multiple resource and consumer species.

### Evolutionary consequences

Besides being crucial to ecology, density dependence and resulting density-dependent selection represent a central link between ecology and evolution (Travis et al., 2013), and is essential for the occurrence of eco-evolutionary feedbacks (Govaert et al., 2019). The shape of density dependence is also crucial for understanding the consequences of adaptive prey evolution (Abrams, 2009a), which is a central topic in eco-evolutionary dynamics research (Yoshida et al., 2003; Hiltunen et al., 2014).

From an evolutionary point of view, Matessi and Gatto (1984) discuss that density-dependent selection (“*K*-selection”) should minimize equilibrium resource availability rather than maximizing 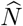 (note parallels to *R*\* theory, Tilman 1980 and MacArthur’s minimum principle, MacArthur 1969, Gatto 1990 and Ghedini et al. 2018). As a consequence, Matessi and Gatto (1984) show that, at the consumer level, 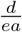 should be minimized by evolution, as this allows to reduce resource availability due to high values of *e* and/or *a* and low values of *d*. Therefore, density-dependent selection can be predicted to increase *r*_0_ (see e.g., Eq. 9) and *β*, either leading to an increase or a decrease in equilibrium densities, depending on resource and consumer behaviour (see Fig. 8). Most importantly, a negative relation does not result from a trade-off between population growth rates and competitive ability, as both parameters are always positively related. The decrease in equilibrium densities is rather a consequence of increased competition. This is an important distinction to classical *r – K* selection assumptions.

**Figure 8:**
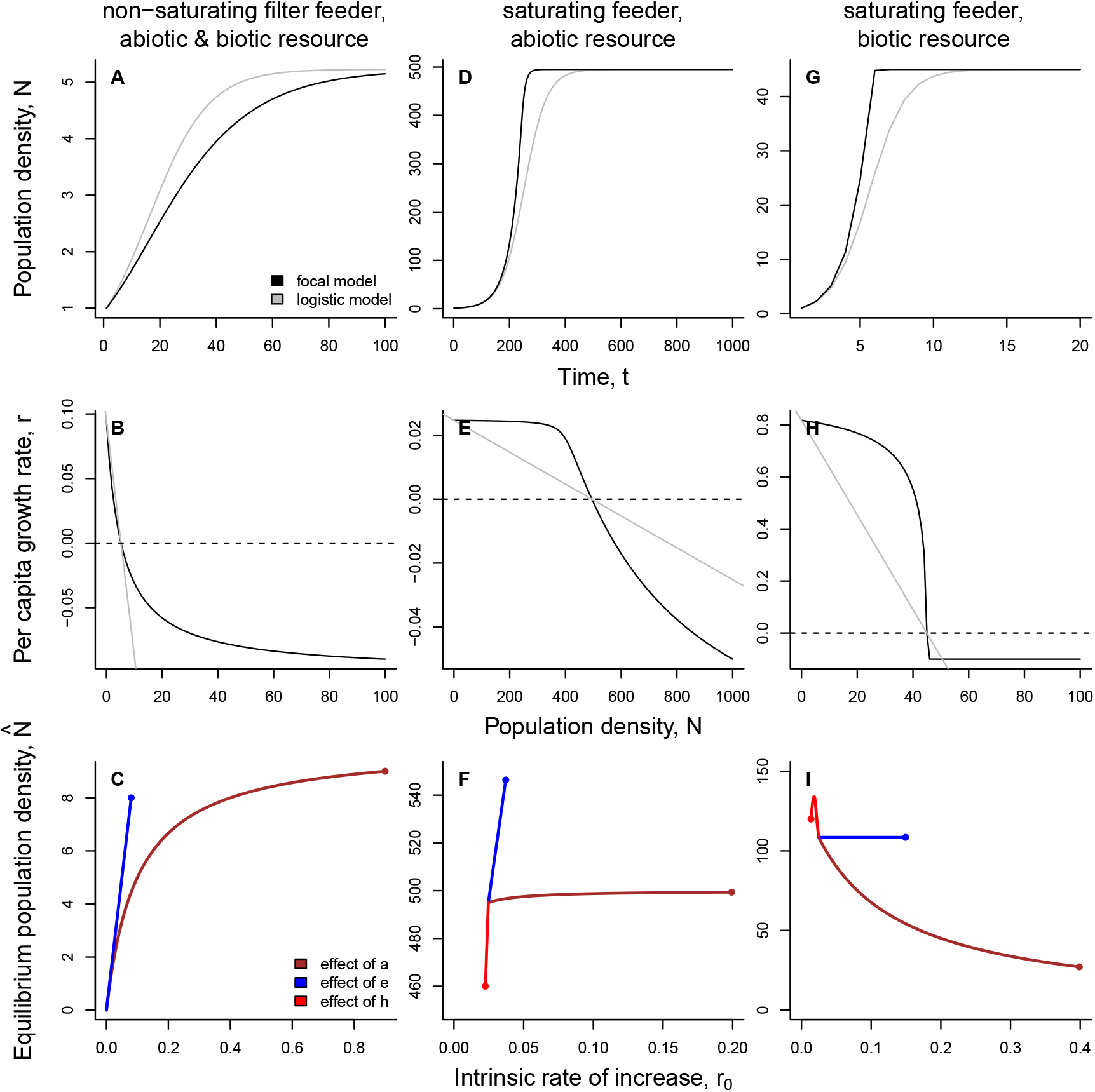
Summary of population growth models (upper row; grey is the logistic for reference), density-regulation functions (central row) and relationships between equilibrium population density 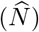 and intrinsic rate of increase (*r*_0_; bottom row). The points in the latter plots visualize the largest *a, e* or *h* value. For more detail see Figs. 2, 4, 6.

Interestingly, Getz (1993) derives *r-K* models from underlying consumer-resource models and shows that *r*_0_ and *K* may be independent if one considers a parameter capturing the maximal growth rate. This parameter only acts on *r*_0_ and may lead to a difference between *r* and *K* selected populations. Some parallels can be found in our work: for both non-saturating filter and saturating feeders, we find that for vanishingly small density-independent mortalities (*d* → 0), the equilibrium population density 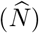 is independent ofthe assimilationcoefficient (*e*). Bycontrast, ifwe assume abiotic resources and vanishingly small density-independent mortalities, the equilibrium population density becomes independent of the maximal foraging rate (*a*). Only in these cases evolution in the respective parameters (*a* or *e*) can lead to a change in growth rate (*r*_0_) without affecting the equilibrium population density 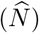.

In a classical *r-K* selection study, Luckinbill (1979) investigated the consequences of adaptation to a low-density environment. Using protists as model organisms, he showed that, in contrast to *r-K* selection theory, *r*-selection did not only lead to an increase in *r*_0_ but simultaneously to an increase in equilibrium densities 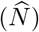. This result can be explained by our work if the protists exhibit a linear functional response (Fig. 8A–C) or feed on abiotic resources (chemostat; Fig. 8D–F). Adaptation to high-density environments may be mainly driven by changes in foraging rates (*a*) as has been prominently investigated in *Drosophila* (Mueller et al., 1991). In line with the prediction of Matessi and Gatto (1984) that selection in high-density environments should minimize 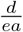, Joshi and Mueller (1988) and Mueller (1990) report that selection in high-density environments increases feeding rate (bite size). Furthermore, Joshi and Mueller (1996) find a trade-off between foraging rate (*a*) and assimilation efficiency (*e*) in *Drosophila*. Similarly, Palkovacs et al. (2011) report that Trinidadian guppies from low-predation environments which have experienced high population densities show adaptations towards increased resource consumption rates.

Most recently, Abrams (2019) has clarified that, in contrast to widely held beliefs, especially in the eco-evolutionary dynamics literature (Hendry, 2017), that adaptation in the consumer usually leads to increases in (equilibrium) population density, maladaptive population density declines can be predicted to occur regularly. In our work, adaptive population density decline should occur especially in consumer-resource systems characterized by biotic resources and saturating consumers (unimodal 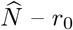 relationships; Fig. 6). Of course it should be noted that such declines may also happen in the other systems considered here: in Figs. 2, 4 and 6 we have only considered that maximally two consumer parameter may change at once, this of course must to be the case. Additionally, the picture may be even more complicated by trade-offs or relationships between the underlying consumer parameters.

Finally, on a larger geographical scale, a widely held assumption is that organisms should be most abundant and exhibit the highest densities in their range core (Burton et al., 2010). However, the generality of this pattern has recently come under scrutiny: in an experimental evolution study, Fronhofer and Altermatt (2015) showed the evolution of lower equilibrium densities in range core populations (see also Fronhofer et al., 2017). More generally, Dallas et al. (2017) could not find increased densities in range cores across 1400 species, including vertebrates and trees. Our current work provides a potential explanation for the lack of this pattern, and even for an inverse pattern: if species feed on biotic resources, exhibit non-linear functional responses, and evolution maximizes foraging rate (*a*) rather than assimilation efficiency (*e*), low equilibrium densities will be the result (Fig. S6).

### Temperature-dependence of population dynamics

Besides having evolutionary consequences, our work has implications for instance in the context of global change research (for a concrete example see Supplementary Material S9). Linking consumer-resource parameters and population level entities explicitly is at the centre of efforts to understand how populations behave under changing climatic conditions, specifically changing temperatures (Uszko et al., 2017). Equilibrium density relates population dynamics to community and ecosystem processes, because it provides a mechanistic link between individual performance, intraspecific interactions, and the total biomass and productivity of a species. Yet, as Gilbert et al. (2014) state, understanding the temperature-dependence of the equilibrium density has long remained a knowledge gap (see also Bernhardt et al., 2018). Important progress has been made recently by Uszko et al. (2017) who for the first time include a mechanistic formulation of prey equilibrium density as a function of temperature in a predator-prey model.

### Multi-species models

While all our considerations have up to now been focused on one consumer species, natural systems rather consist of communities of multiple consumers and resources. Based on our derivations, one can expand our work to multiple species (for some examples, see Supplementary Material S10). For example, Abrams (2009c) presents an extension of Abrams (2009b) to include multiple resource species (see also O’Dwyer, 2018). Extensions to include more complex trophic and non-trophic interactions via additional mortality terms are also possible. Of course, if such complexities are relevant to a biological system, it may be useful to model these interactions explicitly using an appropriate foodweb model instead of attempting a simplification by focusing on the consumer level. This would also resolve intrinsic weaknesses of our approach due to the issues associated with the time-scale separation argument (O’Dwyer, 2018). However, our models remain relevant for example for describing the lowest level represented in such a food web. The shape of resource density dependence has for instance been shown to be crucial when understanding the consequences of adaptive prey evolution (Abrams, 2009a).

### Discussion and conclusion

Our results, based on deriving density-regulation functions from underlying consumer-resource models, show that the parameters of population level growth models (e.g., *r*_0_ and *β*) and equilibrium population densities 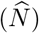 are not independent, but are all functions of consumer traits such as foraging rate and assimilation efficiency (Figs. 2, 4, 6 and 8). This is in good accordance with Matessi and Gatto (1984) and holds true regardless of the underlying consumer-resource dynamics. In contrast to previous work, we derive a priori expectations about the relationship between growth rates, competitive abilities and equilibrium densities that capture the essence of three different consumer-resource systems: non-saturating filter feeders, saturating feeders and consumers feeding on abiotic resources (Fig. 8). Contrary to the widely held belief that growth rates and equilibrium densities are negatively related, we show that various relationships must be expected, depending on the underlying consumer-resource dynamics (Fig. 8). In accordance with previous work (e.g., Thieme, 2003; Johst et al., 2008; Abrams, 2009b; Reynolds and Brassil, 2013) we show that (i) the logistic model (Eqs. 1 and 2) assuming linear density dependence may only be appropriate under very specific conditions, such as competition for spatial resources like nesting sites or territories (see Supplementary Material S1; see also O’Dwyer 2018), and that (ii) most ecological systems will rather follow concave or convex density-regulation functions (Fig. 8), because most systems are bottom-up regulated (Begon et al., 2006). As discussed in Abrams (2009b), the curvature of these density-regulation functions is different from what the θ-logistic model (Gilpin and Ayala, 1973) can achieve. We here show that non-saturating filter feeders, and generally organisms with linear functional responses (Jeschke et al., 2004), follow a convex density-regulation function that is exactly described by a continuous-time version of the Beverton-Holt model (Eq. 8; see also Thieme 2003 and Pástor et al. 2016, for example). For consumers feeding on abiotic resource and following a saturating functional response, we provide a mechanistically derived density-regulation function (Eq. 11). More complex consumer-resource systems can be approximately described by a continuous-time version of the Maynard Smith-Slatkin model (Eq. 13).

Based on our theoretical work, we predict non-saturating filter feeders to generally exhibit a positive relationship between growth rates, competitive abilities and equilibrium population densities (Fig. 2 and 8A–C). The form of this relationship will depend on whether higher growth rates are achieved by increasing foraging rates or by increasing assimilation efficiencies (Fig. 2 and 8C). For foraging strategies that are characterized by a type II, that is, saturating functional response (Fig. 8D–F) while keeping the resources abiotic, we show that the appropriate density-regulation function can be both concave and convex (Fig. 3 and 8E). Increasing foraging rates and assimilation coefficients will still increase both growth rates and equilibrium densities (Fig. 4 and 8F), while the effect of the half-saturation constant is the opposite. The relationship between growth rates and equilibrium densities may be different for more complex systems characterized by both biotic resources and non-linear functional responses (Fig. 8G–I). Specifically, changing foraging rates and half-saturation constants may lead to non-monotonic relationships between growth rates and equilibrium densities (Fig. 6). These considerations highlight that population growth and competitive ability are related, usually positively, and that equilibrium densities are a result of underlying consumer-resource dynamics that may change positively or negatively with population growth and competitive ability. This has important implications for evolutionary considerations as discussed above. Recent microbial work has started to explore the underlying genetic basis of *r – K* relationships (Wei and Zhang, 2019).

Previous work has investigated the shape of density-regulation functions empirically, by using times-series data from growth experiments and natural population dynamics. For example, Borlestean et al. (2015) investigated the curvature of density dependence in a *Chlamydomonas* chemostat. While the authors report to be surprised by the general convexity of the density-regulation function, these results are in perfect accordance with our predictions (Fig. 8B). In a comparative study, Sibly et al. (2005) used a large dataset to show that the relationship between growth rate and density is generally convex, that is, exhibits first a fast decrease and then slows down (*θ* < 1 in the *θ*-logistic model). This result corresponds to our scenario for non-saturating filter feeders and abiotic resources (Fig. 8B), which is rather surprising, given that the dataset included mammals, birds, fish and insects. However, as Clark et al. (2010) laid out in detail, the dominance of *θ* < 1 values may be due to technical difficulties in fitting the *θ*-logistic model. Nevertheless, this study, along with the findings of Eberhardt et al. (2008) who suggested that the *θ*-logistic usually outperforms the logistic in a number of vertebrate species, clearly highlight the general non-linearity of density-regulation functions. These examples show the value of our work, as it provides theoreticians and empiricists with theoretically grounded assumptions on the occurrence of specific shapes of density dependence and trait relationships. Besides the implications discussed above, the curvature of density dependence itself is highly relevant for a population’s response to stressors. Hodgson et al. (2017) demonstrated that when density dependence is concave the effect of stressors on focal populations is always amplified, while the response can be amplified or dampened for convex density dependencies. For a detailed discussion of the impact of the shape of density dependence in basic and applied ecology see Abrams (2009c).

Of course, it is important to keep in mind that our models are simplified representations of population dynamics and ignore, for instance, age or stage structure (Mueller, 1997). We do not include Allee effects (Allee, 1931; Courchamp et al., 2008), that is, reduced population growth at low densities. There are numerous mechanisms that can lead to Allee effects and we speculate that these may lead to different functional relationships between population growth and population density. At a descriptive level, demographic Allee effects can be included as shown in Kubisch et al. (2016) for discrete time systems, for example. Our considerations also do not include time-lags in density dependence (Ratikainen et al., 2008) which are relevant in both theoretical and applied contexts, for example for population stability. For reasons of space we also do not discuss discrete time models. Note that Turchin (2003) and Thieme (2003) treat this topic in detail and mechanistic derivations of the Ricker model have, for example, been used by Melbourne and Hastings (2009).

Finally, we would like to reiterate the point made by Mallet (2012): we cover a long-discussed topic in ecology. Multiple authors have noted difficulties with the *r – K* formulation of the logistic and potential non-linearities in density dependence. Advanced textbooks like Thieme (2003) and Pástor et al. (2016) have shown that for non-saturating filter feeders and abiotic resources the continuous-time version of the Beverton-Holt model can be derived (see also Abrams, 1977; Schoener, 1978). Mallet (2012) discusses how using the *r – α* formulation of the logistic, where appropriate, alleviates some of the problems encountered with the *r – K* formulation. While these considerations are highly relevant to both empiricists and theoreticians, they seem to remain largely ignored. We hope that the insights provided here, as well as our expansion of past work towards more complex consumer-resource interactions resulting in an understanding of functional relationships between population level parameters, will help to change how density regulation and relationships between equilibrium density and population growth are treated in ecology, evolution and beyond (Aktipis et al., 2013).

## Author contributions

E.A.F. conceived the study. All authors discussed the content and topic of the study. E.A.F., L.G. and S.J.S. analysed the mathematical models. E.A.F. gathered the empirical data. E.A.F. performed the statistical analyses including fitting of population growth models. E.A.F. wrote the manuscript and all authors commented on the draft.

## Acknowledgements

We thank Peter Abrams, François Massol as well as three anonymous referees for valuable comments on the previous versions of the text. F.A. received funding from the Swiss National Science Foundation Grant No PP00P3_I79089 and the University of Zurich Research Priority Programme “URPP Global Change and Biodiversity”. S.J.S. is supported by US NSF DMS-1716803. This is publication ISEM-YYYY-XXX of the Institut des Sciences de l’Evolution – Montpellier.

## Data availability

Data will be deposited in Dryad and R-code will be made available via GitHub and a Zenodo DOI.

## Supplementary Material

### S1 Derivation of the logistic model by Lakin and Van Den Driessche (1977)

Multiple authors have reported that one can derive the logistic growth model (Eqs. 1 or 2) for consumers from underlying consumer-resource dynamics (e.g. MacArthur, 1970; Thieme, 2003; Abrams, 2009; Reynolds and Brassil, 2013). Most of these derivations, however, assume that the underlying resource dynamics (Eq. 3a in the main text) follow a logistic growth model (Eqs. 1 or 2 in the main text) which, given the principle of inheritance of the curvature of density dependence reported by Abrams (2009), appears circular. Reynolds and Brassil (2013) justify their usage of the logistic for describing resource dynamics by referring to work by Lakin and Van Den Driessche (1977) who indeed derive the logistic from a chemostat model. In the Lakin and Van Den Driessche (1977) model resource dynamics (*R*) are described as follows:

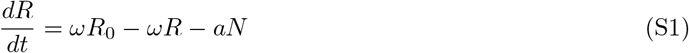

with *ω* being the flow rate into and out of the system, and *R*_0_ the resource concentration flowing into the system. The consumer dynamics (*N*) are thought to follow

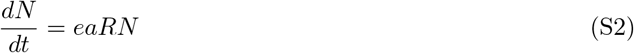

where *e* is the assimilation efficiency of the consumer (*N*) and a its foraging rate. As described in the main text (Eq. 4), one can assume that the resources are at equilibrium 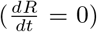 and solve Eq. S1 for *R* to obtain the equilibrium resource population density 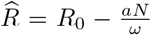. Then the per capita consumer dynamics follow

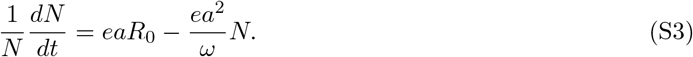

This formulation is identical to the linear density-regulation function assumed in the Verhulst (1838) model (Eq. 2 in the main text) with setting *r*_0_ = *eaR*_0_ and 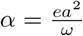 (see also Fig. S1). While this result may seem satisfactory, it is worth noting that in Eq. S1 the consumer harvests a constant amount of resources (*aN*; linear functional response), regardless of resource availability. Biologically, this seems a rather unrealistic assumption.

Despite its artificiality, this model already hints at the non-independence of population growth rates (*r*_0_) and competitive abilities (here *α*, the slope of the density-regulation function). As depicted in Fig. S1 population growth rates and competitive abilities will covary. The relationship is either linear or quadratic, depending on whether the assimilation efficiency (*e*; Fig. S1B) or the foraging rate (*a*; Fig. S1A) is the underlying driver (since r_0_ = *eaR*_0_ and 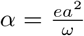). It is also interesting to note that the equilibrium density, here defined as 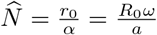 is not a function of the assimilation efficiency (*e*) and decreases with increasing foraging rate (*a*).

**Figure S1:**
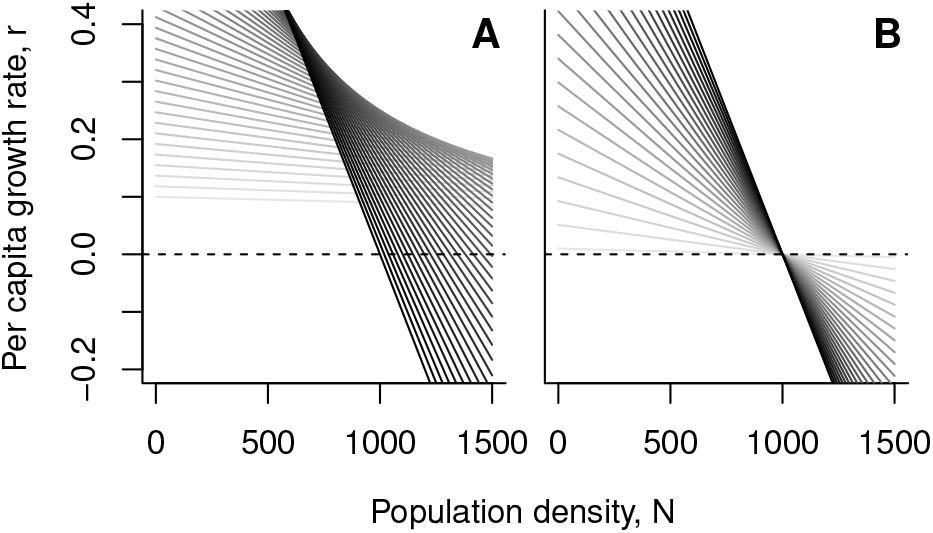
Density dependence for a consumer following the Lakin and Van Den Driessche (1977) model. (A) Effect of changing the foraging rate (*a*) while keeping the assimilation efficiency constant (*e* = 0.01). Darker shades of grey indicate higher parameter values. (B) effect of changing the assimilation efficiency parameter (*e*) while keeping the foraging rate constant (*a* = 0.1). Parameter examples: *R*_0_ = 1000, *ω* = 0.1, *e* ∈ [0.0001, 0.01], *a* ∈ [0.01, 0.1].

### S2 Consumer dynamics with a linear functional response and abiotic resources – an alternative chemostat model

As an alternative to Eq. 5 and following for instance Abrams (1988) we may assume the following relationship which does not include a flow rate:

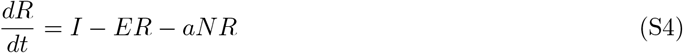

while keeping Eq. 6 for the consumer.

The equilibrium resource concentration is then given by 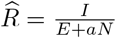. Substituting this into Eq. 6 and rearranging gives us exactly the Beverton-Holt model (Eq. 8) with 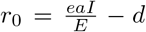 and the competition coefficient 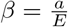. The shape of resulting 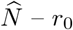 relationships are summarized in Fig. S2.

**Figure S2:**
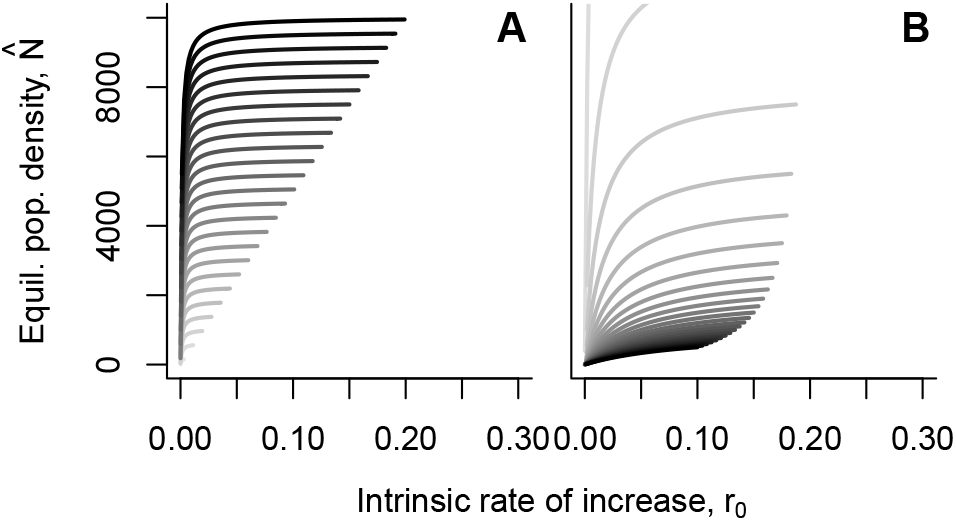
Relationship between intrinsic rate of increase (*r*_0_) and equilibrium population density 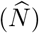 for consumers following a Beverton-Holt density-regulation function (chemostat model given in Eq. S4 for the resource and linear functional response for the consumer). (A) Effect of changing the foraging rate (*a*; solid lines; dots represent the *r*_0_ and 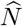 values for the highest value of *a*) and the assimilation efficiency (*e*; darker shades of grey indicate higher values of *e*) while the death rate is kept constant (*d* = 0.001). (B) Effect of changing the foraging rate (*a*; solid lines; dots represent the *r*_0_ and 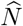 values for the highest value of *a*) and the death rate (*d*; darker shades of grey indicate higher values of *d*) while the assimilation efficiency is kept constant (*e* = 1). Parameter examples: *I* = 10, *E* = 5, *e* ∈ [0.02, 1], *a* ∈ [0.0001, 0.1], *d* ∈ [0, 0.1].

### S3 Consumer dynamics with a type II functional response and abiotic resources

For a consumer (*N*) with a type II functional response feeding on an abiotic resource (*R*; chemostat model) we assume the following dynamics:

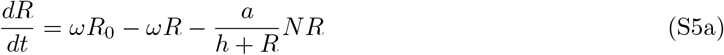

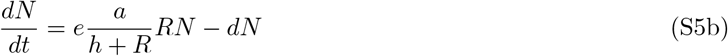

For any consumer density (*N*), the growth rate 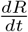 of the resource is a continuous, decreasing function of *R* and takes on the value *ωR*_0_ > 0 at *R* = 0 and becomes negative for sufficiently large *R*. Thus, there is a unique positive value 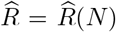 at which 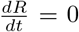 for the given consumer density *N*. When the resource dynamics occurs at a much faster time scale than the consumer dynamics (see, e.g, Rinaldi and Muratori, 1992; Hek, 2009), this resource density 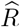 corresponds to a quasi-steady state. To find this quasi-steady state, we have to solve the following equation:

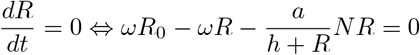

Assuming that *h* + *R* ≠ 0, we can multiply both sides with *h* + *R*. Rearranging terms results in the following quadratic equation:

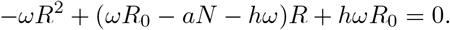

This quadratic equation has two solutions:

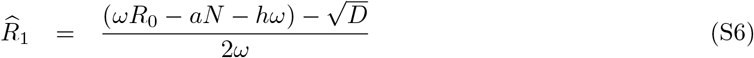

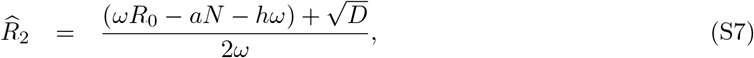

with *D* = (*ωR*_0_ – *aN*)^2^ + 2*hω*(*ωR*_0_ + *aN*) + *h*^2^*ω*^2^. By uniqueness and positivity of 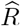, we have 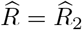. Substituting 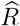 into Eq. S5b results in

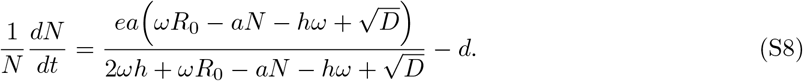

Theorem 1 of Schreiber (1996) implies that the consumer-resource system (S5a)–(S5b) always has a unique globally stable equilibrium – in particular, there are no consumer-resource oscillations. Consequently, the one-dimensional system (S8) provides a reasonable approximation of the consumer dynamics on the slower time scale for all parameter values and all initial conditions.

Equation (S8) is of the following form

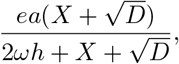

with *X* = *ωR*_0_ – *aN* – *hω*. We can then use the binomial product (*a* + *b*)(*a* – *b*) = *a*^2^ – *b*^2^ to simplify this equation by multiplying the numerator and denominator with 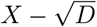. This results in:

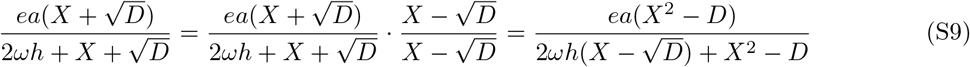

where 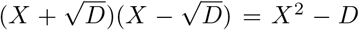. Substituting *X = ωR*_0_ – *aN – hω* and *D* = (*ωR*_0_ – *aN*)^2^ + 2*hω*(*ωR*0 + *aN*) + *h*^2^*ω*^2^ into *X*^2^ – *D* gives

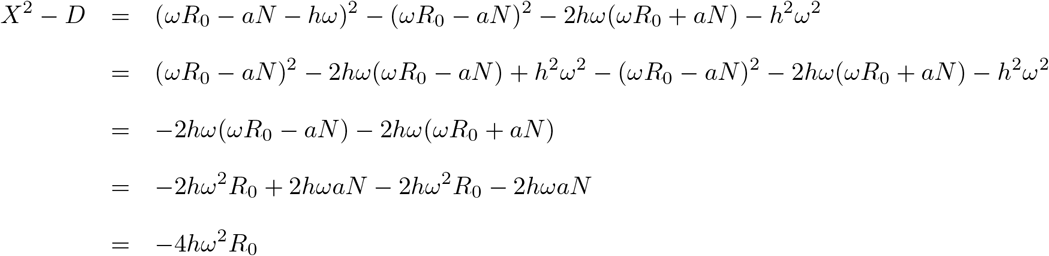

For the denominator we get 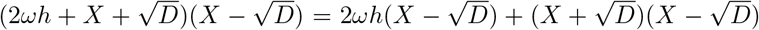. For the second term we already know that it equals to –4*hω*^2^*R*_0_. Thus the denominator becomes:

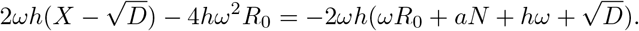

We thus simplify Eq. S8 to:

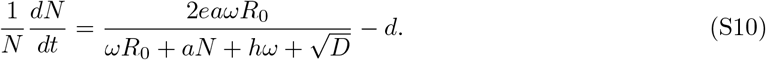

Now we can solve Eq. S10 to find the density *N* at equilibrium 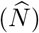. We first need to assume that 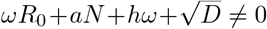. Note that we can rewrite Eq. S10 to an expression of the form 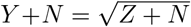 by rearranging Eq. S10 to

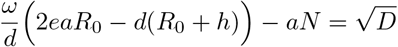

Now squaring both sides and solving to *N* results in:

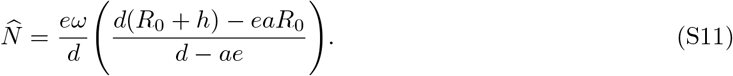

From Eq. S10 we can calculate *r*_0_ by assuming *N* = 0:

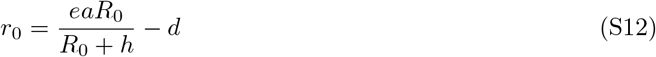

From this we find that

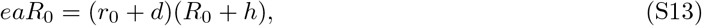

which we can substitute into Eq. S10. This then gives the density-regulation function for consumer dynamics with a type II functional response and abiotic resources:

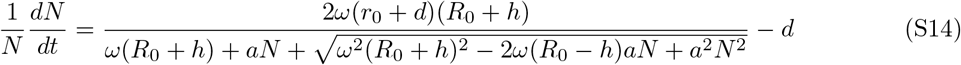

We can simplify this equation by defining the competitive ability *β* as in Eq. 8 as 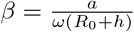:

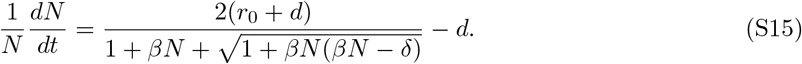

with 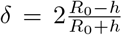. If *δ* → 2, then the denominator of Eq. S15 simplifies to 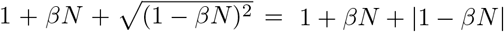. Hence, when 1 – *βN* ≥ 0, this simplifies Eq. S15 to

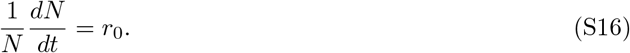

This implies exponential growth as a result of *h* → 0, that is, foraging being resource independent. By contrast, if *δ* → –2, then the denominator of Eq. S15 simplifies to 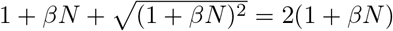. This reduces Eq. S15 to

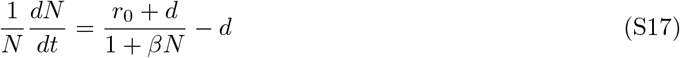

which results into the Beverton-Holt model (Eq. 8 in main text). This result is expected, because *δ* → –2 is a result of *h* → ∞ which leads to a linear functional response. Solving equation (S15) to equilibrium density 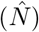 then gives:

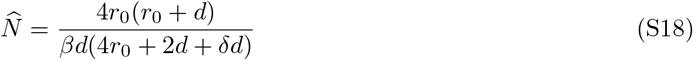

### S4 Curvature of the density-regulation function – type II functional response and abiotic resources

We will use the following general consumer-resource model

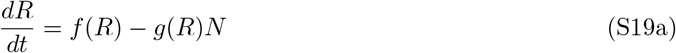

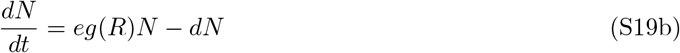

where *R* is the resource and *N* the consumer population density. The function *f*(*R*) captures the growth of the resources and the function *g*(*R*) captures the functional response of the consumer. In order to have a general idea of the form of the per-capita growth function of the consumer 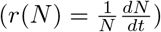, we can take the first and second derivative with respect to *N* of *r*(*N*). This then equals:

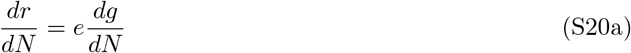

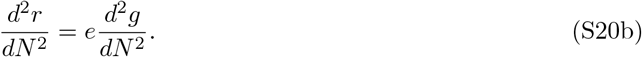

Hence, if we have information on the sign of 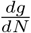 and 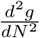, this will provide information on the sign of 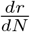 and 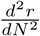. But as *g* is a function of *R* and *R* is a function of *N*, the first and second derivative of *g* to *N* can be rewritten as:

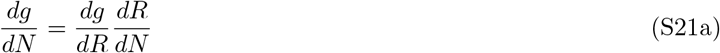

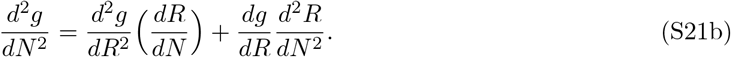

So we need to find expressions for 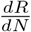 and 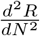. To do so, note that at resource equilibrium 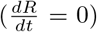, we find that

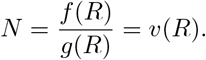

If we set this fraction to a function *v*(*R*), and implicitly differentiate to *N*, this yields:

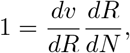

as *R* is a function of *N*. From which it follows that

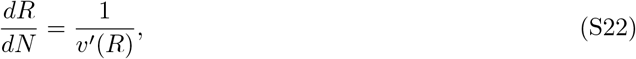

where 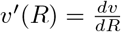. A second implicit differentiation yields:

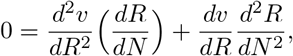

which gives after rearranging terms

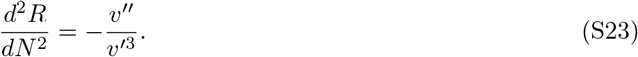

Eq. S23 determines the curvature of *R* as a function of *N*. Hence, Eq. S22 and Eq. S23 gives an expression for the first and second derivative of *R* in function of *N*, which we can use to substitute in the first and second derivative of *g* to *N*:

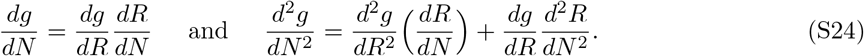

For abiotic resources with type II functional response, function *f* and *g* become:

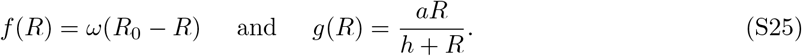

In this case, v(R) and its first and second derivative equal:

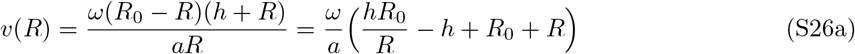

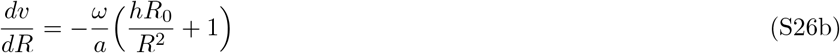

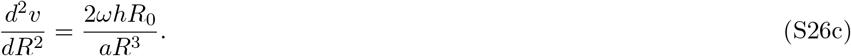

From this it follows that 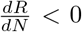 and 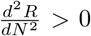. Thus, the resource equilibrium density is a decreasing convex function of consumer population density, just as in the case of the linear functional response.

From this we can determine the curvature of the per-capita growth rate by substituting the information given in Eq. S25 into Eq. S24. We get:

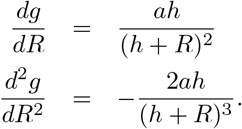

From this Eq. S24 becomes

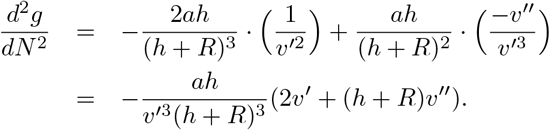

As *v*’ < 0, it follows that 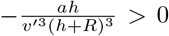. Hence 2*v*’ + (*h* + *R*)*v*″ determines the sign of 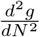. Using information from Eq. S26, we can replace *v*’ and *v*″ in 2*v*’ + (*h* + *R*)*v*″ and obtain:

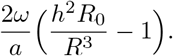

Now 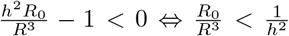. Hence, *R* decreases with *N* from *R*_0_ at *N* = 0 until *N* is such that *R*^3^ = *h*^2^*R*_0_. At *N* = 0, *R* = *R*_0_, and we find that 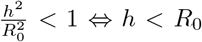. Note that the concavity in the density-regulation function stems from the concavity of the type II functional response of the consumer, not from the resource level itself, as the resource is a decreasing convex function of consumer population density.

### S5 Fitting known density-regulation models to consumer dynamics with a type II functional response and abiotic resources

In order to compare the behaviour of Eq. S15 to existing population-regulation models, we take the same approach as Abrams (2009) and fitted these models to Eq. S15 using a least-squares approach. The respective ODEs were solved (function ‘ode’ of the ‘deSolve’ package in R version 3.4.4) and the model was fitted by minimizing the sum of squared residuals (Levenberg-Marquardt algorithm implemented in function ‘nls.lm’ of the ‘minpack.lm’ package in R). Specifically, we here consider the Beverton-Holt model (Eq. 8), the *θ*-logistic model (Gilpin and Ayala, 1973), the Hassel model (Hassell et al., 1976) and the Maynard Smith-Slatkin model (Maynard Smith and Slatkin, 1973) as possible candidates.

**Figure S3:**
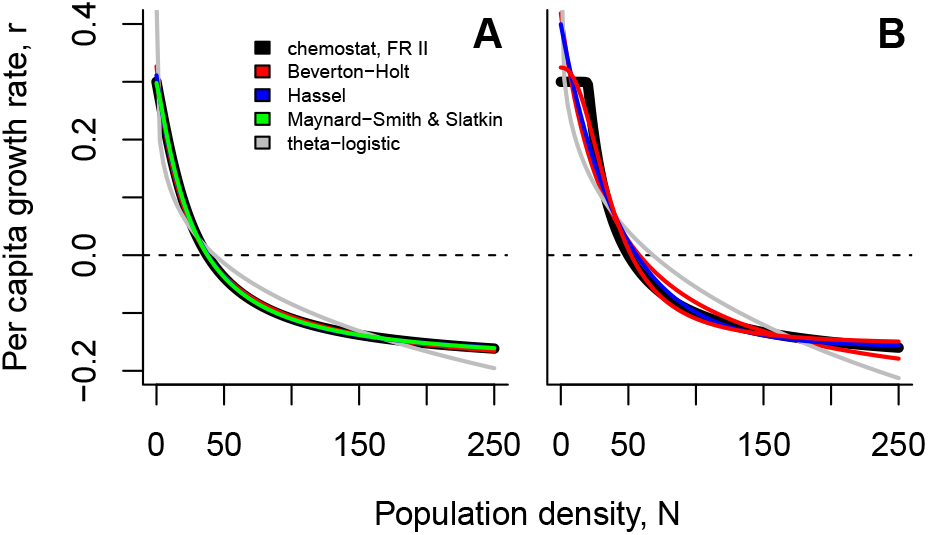
Fitting existing population growth models to the population growth function of consumer with a type II functional response feeding on abiotic resources (chemostat model). (A) High half-saturation constant and foraging rate (*h* = 1000, *a* = 10) (B) Low half-saturation constant and foraging rate (*h* = 1, *a* = 5). Parameter examples: *R*_0_ = 100000, *ω* = 0.1, *e* = 0.1, *d* = 0.2.

Because the Maynard Smith-Slatkin model approximates the dynamics of Eq. S15 sufficiently well, we can use this model to analyse numerically how changes in consumer-resource parameters impact the parameters used in the Maynard Smith-Slatkin model (Fig. S4). We recapture the results laid out in the main text and show that strong concavity (determined by the shape paramete*r γ*) mainly occurs at low values of the half-saturation constant (*h*) and specifically for low foraging rates (*a*).

**Figure S4:**
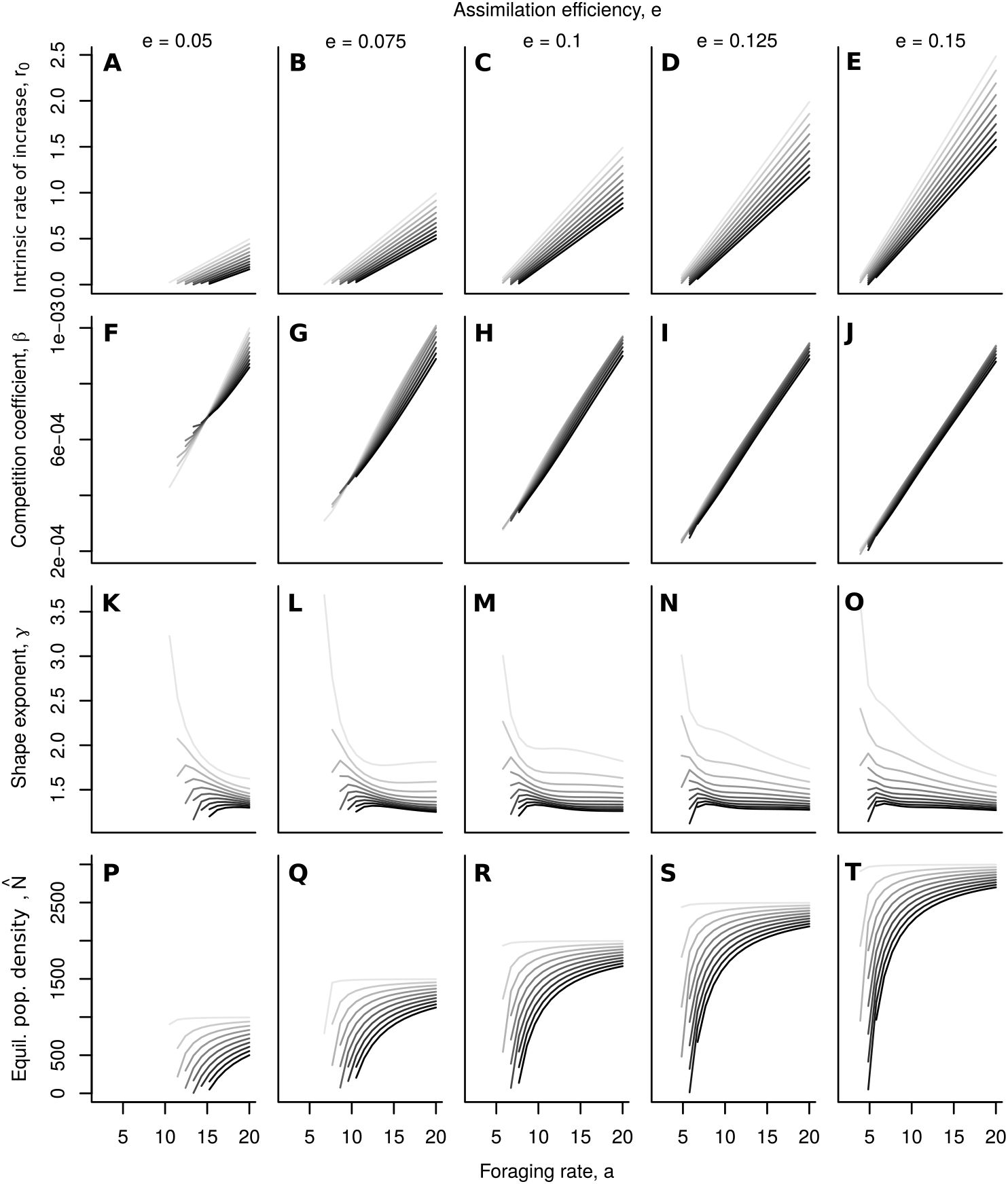
Relationships between consumer-resource parameters and the parameters of the Maynard Smith-Slatkin model for a consumer with a type II functional response feeding on abiotic resources. The Maynard Smith-Slatkin model was fitted using a least-squares approach to the respective realisation of Eq. S15. Darker shades of grey indicate higher parameter values of the half-saturation constant (*h* ∈ [500, 50000]). The assimilation efficiency (*e* ∈ [0.05, 0.15]) increases from left to right. Parameter examples: *a* ∈ [2, 20], *R*_0_ = 100000, *ω* = 0.1, *d* = 0.5.

### S6 Consumer dynamics with a type II functional response and biotic resources

For a consumer-resource system with biotic resources (Eq. 8) and a consumer exhibiting a type II functional response, we can write:

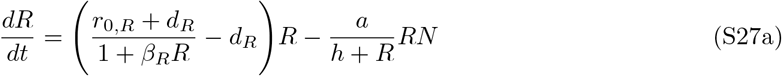

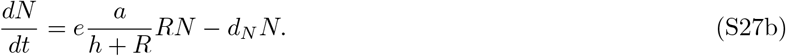

Consumer-resource dynamics of this type have been studied extensively (see, e.g., Cheng et al., 1982; Kuang and Freedman, 1988). Let *α* = *r*_0_,*R*+ *d_R_*. The consumer and resource can coexist, in the sense of permanence (Schreiber, 2000), if and only if *α* > *d_R_, ea* > *d_N_*, and 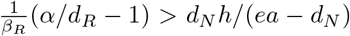. When this coexistence occurs there is a unique equilibrium supporting both species. The stability of this equilibrium depends on the slope ofthe non-trivial resource nullcline (given by *N*= (*h+R*)(*α*/(1+*β_R_*)– *d_R_*)/*a*) at the point at which it intersects the non-trivial consumer nullcline (given by *R* = *d_N_h*/(*ea–d_N_*). Iftheslope isnegative, then Cheng et al. (1982) prove the consumer-resource equilibrium is globally stable. If the slope is positive, then Kuang and Freedman (1988) prove that the equilibrium is unstable and there is a unique limit cycle at which the consumer and resource ultimately coexist.

If we assume that the resource dynamics occur at a faster time scale then the consumer dynamics, then we can solve for potential quasi-steady states for the resource at any consumer density *N*. Unlike the case of the abiotic resource dynamics, there does not always exist a unique, positive resource quasi-steady state. To understand when such a state does or does not exist, we need to solve 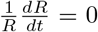 for *N* which gives the non-trivial resource nullcline

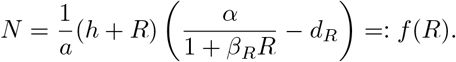

Whether or not *f* (*R*) for *R* ≥ 0 is hump-shaped or just decreasing in *R*, depends on *f*’(0) = (*α* – *d_R_* – *αβ_R_h*)/*a* is positive or negative. If *f* ‘(0) ≤ 0, *f* (*R*) decreases for all *N* and the maximum value of *f* (*R*) is *N_max_* = *f*(0) = *h*(*α* – *d_R_*)/*a*. If *f*’(0) > 0, then there is a critical value 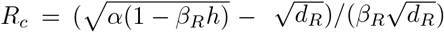 at which *f*’(*R_c_*) = 0 in which case the maximum value of *f*(*R*) is *N_max_* = *f*(*R_c_*).

If *N* > *N_max_*, solving *dR/dt* = 0 for the quasi-steady value of *R* always yields *R* = 0 as the only nonnegative solution. Hence, for these values of the consumer density, the consumer is driving the resource extinct. If *N* < *N_max_*, there will be at least one positive solution to solving *dR/dt* = 0 for *R*. Solving this equation yields two roots of a quadratic equation:

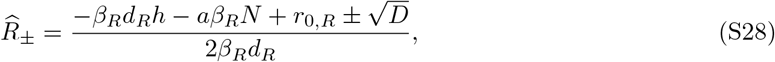

with 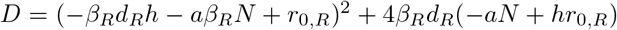. If *f*’(0) ≤ 0 and *N* < *N_max_*, then 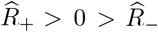. Hence, in the fast-slow limit, the resource density quickly approaches the quasi-steady state 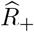. The same conclusion applies when *f*’(0) > 0 and *N* ≤ *f* (0). When *f*’(0) > 0 and *f* (0) < *N* < *N_max_*, both quasi-steady states 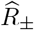 are positive. However, in the fast-slow limit, the resource density quickly approaches the quasi-steady state 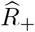 whenever its initial density is 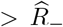 and quickly approaches 0 otherwise. 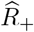 are stable at the fast-time scale, while 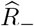 is unstable at this time scale.

Putting all of these results together, we have the following approximation for the consumer dynamics:

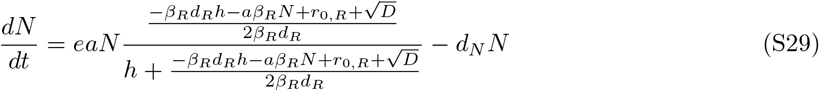

However, this approximation is only valid as long as *N* < *N_max_*. When the equilibrium of the full system (S27a)–(S27b) is globally stable, this constraint holds for all *t* providing that *N*(0) < *N_max_*. Alternatively, when the equilibrium of the full system (S27a)–(S27b) is unstable, solutions *N*(*t*) of (S29) with *N*(0) < *N_max_* will in finite time increase to value *N_max_* at which time the approximation is no longer valid.

Using 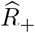, we can simplify Eq. S29 using similar steps as in Supplementary Material S3. Setting *X* = – *β_R_d_R_h* – *β_R_aN*+*r_0,R_*, Eq. S29 can be rewritten as follows:

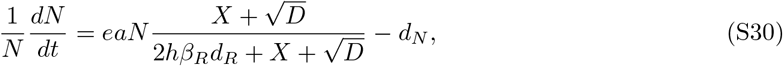

Multiplying the numerator and denominator with 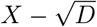 and simplifying gives:

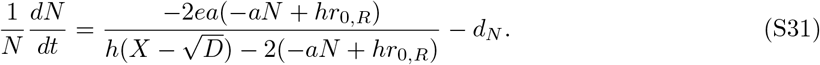

In order to find the equilibrium density 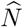, we can solve Eq. S31 for *N*, giving the solutions: 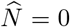 and

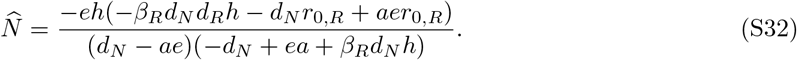

*r*_0_ can be obtained by setting *N* = 0 in Eq. S31, which gives:

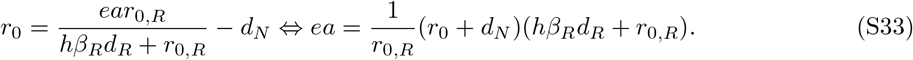

Substituting *r*_0_ into 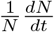 by using Eq. S33 givesg

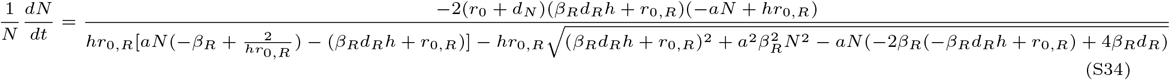

The previous equation can then be simplified by dividing by *β_R_d_R_h* + *r*_0_,*R*:

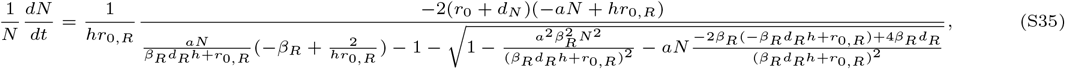

and by setting 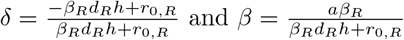 and 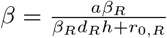 we can rewrite the previous equation as follows:

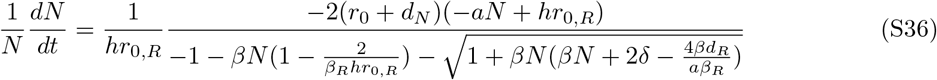

which equals

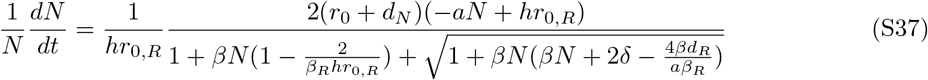

### S7 Fitting known density-regulation models to consumer dynamics with a type II functional response and biotic resources

For details of the fitting procedure see Supplementary Material S5.

**Figure S5:**
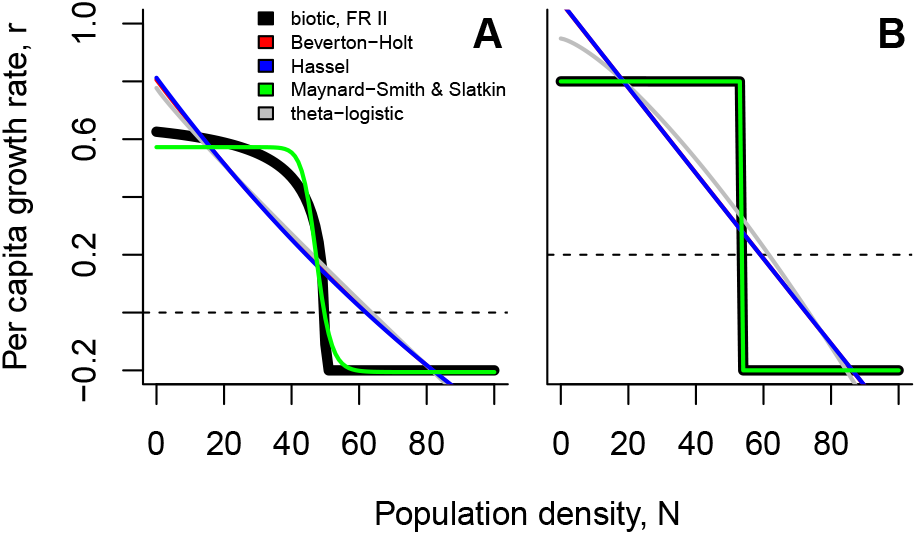
Fitting existing population growth models to the population growth function of consumer with a type II functional response feeding on biotic resources (Beverton-Holt model; Eq. 8). (A) High half-saturation constant and foraging rate (*h* = 900, *a* = 9) (B) Low half-saturation constant and foraging rate (*h* = 1, *a* = 5). Note that blue (Hassel model) and red lines (Beverton-Holt model) overlap completely here. Parameter examples: *r_0,R_* = 0.5, *β_R_* = 0.001, *d_R_* = 0.05, *e* = 0.1, *d* = 0.2.

Fitting the Maynard Smith-Slatkin model to a wide range of parameter combinations highlights that relations between intrinsic rates of increase (*r*_0_) and competition coefficients do not change qualitatively, while relations with the equilibrium density do. Up to here, for abiotic or biotic resources and non-saturating filter feeding consumers, the equilibrium density was always a monotonically increasing function of foraging rates. We here find that this relationship is unimodal or monotonically decreasing for biotic resources and saturating consumers (Fig. S6). The abrupt changes from concave to convex of the density-regulation function (see Fig. 5) lead to high values of the shape parameter (*γ* 1) of the fitted Maynard Smith-Slatkin model (Fig. S6 K–O).

**Figure S6:**
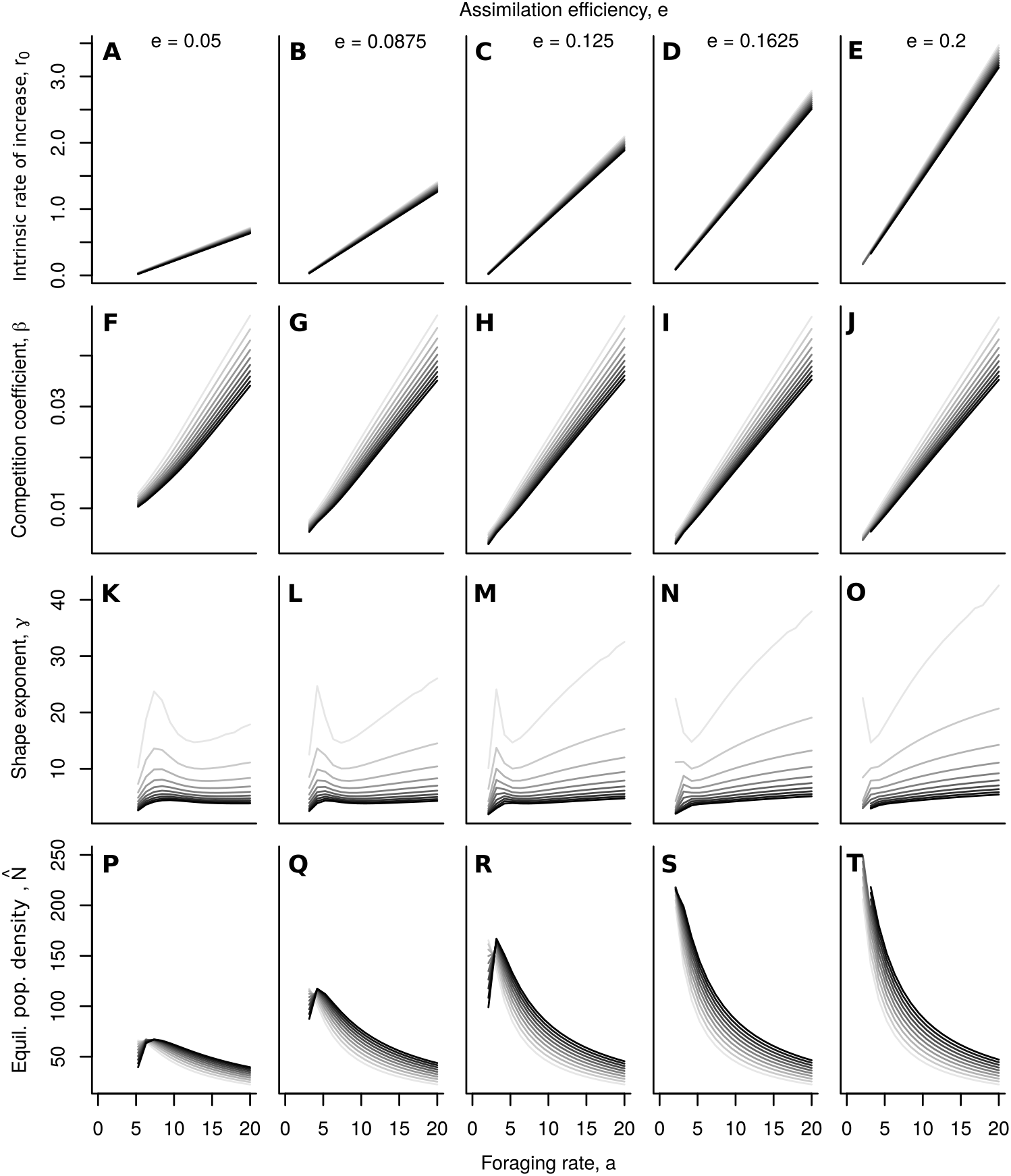
Relationships between consumer-resource parameters and the parameters of the Maynard Smith-Slatkin model for a saturating consumer (type II functional response) feeding on biotic resources (following the Beverton-Holt model). The Maynard Smith-Slatkin model was fitted using a least-squares approach to the respective realisation of Eq. S37. Darker shades of grey indicate higher parameter values of the half-saturation constant (*h* ∈ [900,2000]). The assimilation efficiency (*e* ∈ [0.05,0.2]) increases from left to right. Parameter examples: *a* ∈ [0,20], *d* = 0.2, *r_0,R_*= 0.5, *β_R_*= 0.001, *d_R_*= 0.05.

### S8 Fitting population growth models to data

#### Study organism, experimental procedure and data collection

We used 7 strains of the freshwater ciliate *Tetrahymena thermophila* (strains: SB3539, B2086.2, A*III, CU438.1, A*V, CU427.4, CU428.2) originally obtained from the Cornell Tetrahymena Stock Center. These strains were kept for maintenance and during the experiments in protist medium (1 g dried *Lactuca sativa* powder in 1.6 L Volvic water) with *Serratia marcescens* as a food resource.

Growth experiments were carried out in 20 mL vials (Sarstedt) at 20 °C. 6 replicated growth curves of each *Tetrahymena* strain were started with 250 μL of protist culture (4 d old, at equilibrium) in 20 mL bacterized medium (10% *Serratia marcescens* from a 3 d old culture that had reached equilibrium). The growth experiment was followed over the course of 2 weeks and 2 mL of medium was replaced in each microcosm three times per week with freshly bacterized new medium.

Data was collected using video recording and analysis. At regular intervals (twice per day for the first two days, subsequently once per day or once every 2–3 days) the microcosms were sampled and the samples were placed on a counting slide (height: 0.5 mm) under a stereomicroscope (Perfex Pro 10) at a 2-fold magnification. Using a microscope camera (Perfex SC38800) videos were recorded for a total duration of 10s at a rate of 15 frames per second imaging a volume of 31 μL. Videos were subsequently analysed using a customized version of the ‘bemovi’ package (Pennekamp et al., 2015) in R.

#### Model fitting

We used the ‘rstan’ package in R to solve the ODEs and fit the 4 potential deterministic growth models (logistic, Eq. 1; Beverton-Holt, Eq. 8; Eq. 11; Maynard Smith-Slatkin, Eq. 13). We use trajectory matching, that is, we assume pure observation error for simplicity. For a detailed description see Rosenbaum et al. (2019). As in such a trajectory matching fitting exercise the fit depends very strongly on the first population density value (*N*_0_) we alleviate this problem to some degree by estimating N0 as an additional parameter. To facilitate fitting, both data and model parameters (except *δ* in Eq. 11) were log-transformed, and, besides fitting the respective parameters of the ODEs we also fitted the initial density as a free parameter. Priors were chosen based on published data on *Tetrahymena* (Fronhofer and Altermatt, 2015; Fronhofer et al., 2017; Altermatt and Fronhofer, 2018). See below for the rstan model code.

**Figure S7:**
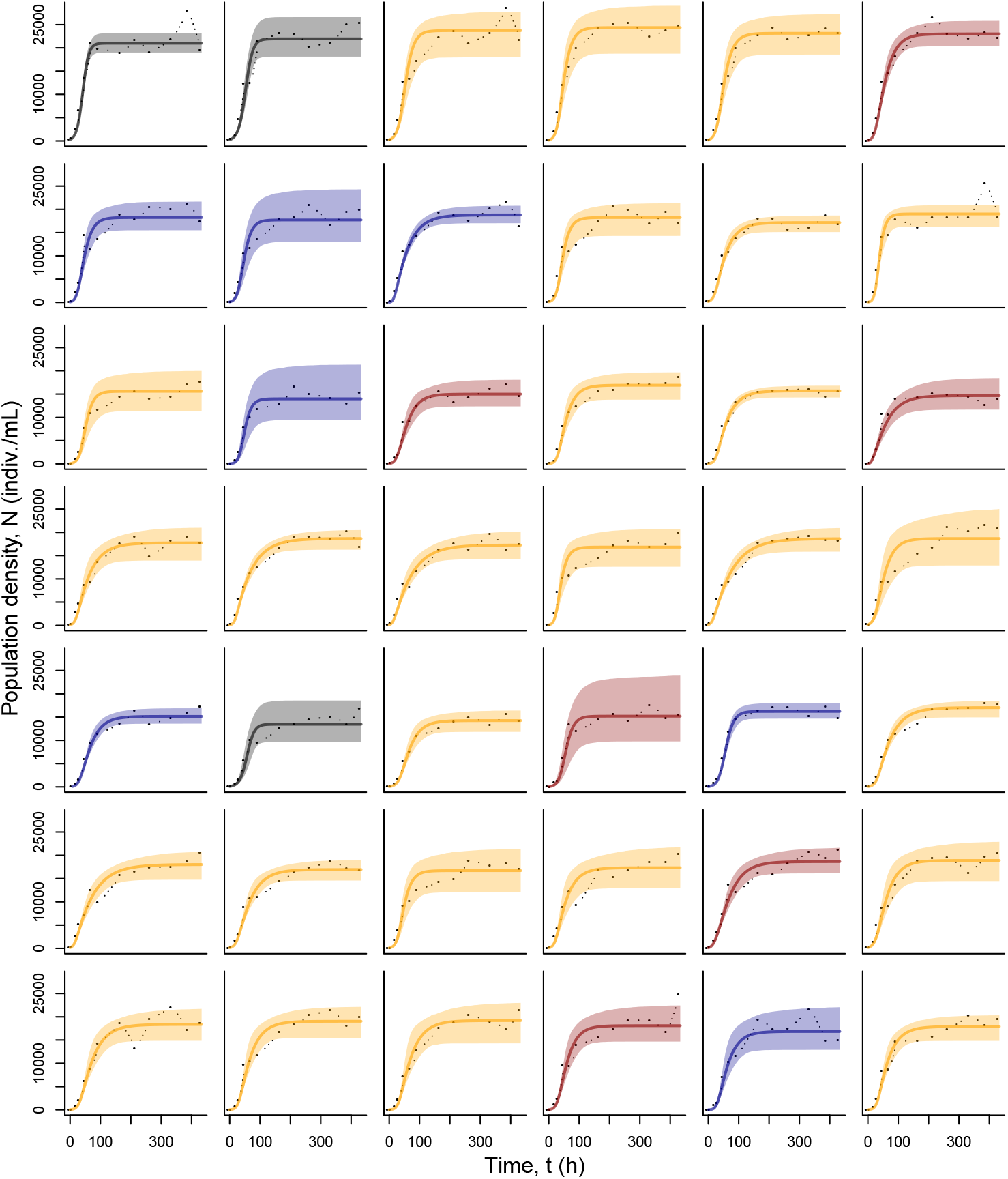
Fitting population growth models to *Tetrahymena thermophila* dynamics using Bayesian inference (see Supplementary Material S8 and Rosenbaum et al. (2019) for details). We fitted the logistic (black; Eq. 1), the Beverton-Holt model (blue; Eq. 8), Eq. 11 (orange) as well as the Maynard Smith-Slatkin model (red; Eq. 13) and compared fits using WAIC (see Tab. S1). Only the best fit is shown. Columns indicate technical replicates of growth curves, while rows show different genotypes of *Tetrahymena thermophila* (strains from top to bottom: SB3539, B2086.2, A*III, CU438.1, A*V, CU427.4, CU428.2).

**Table S1:** Model selection results (WAIC values) for all mating types and replicates visualized in Fig. S7. We fitted the logistic (Eq. 1; “r–K”), the Beverton-Holt model (Eq. 8; “BH”), Eq. 11 as well as the Maynard Smith-Slatkin model (Eq. 13; “MSS”).

| **Mating type** | **Replicate** | **WAIC(r-K)** | **WAIC(BH)** | **WAIC(Eq. 11)** | **WAIC(MSS)** |
| --- | --- | --- | --- | --- | --- |
| I | 1 | -11.32 | -11.28 | -10.66 | -10.89 |
| I | 2 | 3.46 | 3.81 | 3.67 | 4.16 |
| I | 3 | 4.05 | 3.64 | 3.46 | 3.88 |
| I | 4 | 6.51 | 5.21 | 5.01 | 5.12 |
| I | 5 | 3.85 | 2.11 | 1.96 | 2.11 |
| I | 6 | -2.21 | -11.97 | -11.62 | -12.01 |
| II | 1 | -0.05 | -0.65 | 0.02 | -0.13 |
| II | 2 | 15.28 | 15.07 | 15.55 | 15.47 |
| II | 3 | 6.15 | -16.26 | -14.80 | -15.43 |
| II | 4 | 3.06 | 3.51 | 2.39 | 3.32 |
| II | 5 | -4.95 | -11.95 | -15.61 | -13.30 |
| II | 6 | -5.92 | -6.42 | -7.37 | -6.62 |
| III | 1 | 12.68 | 12.66 | 12.22 | 12.71 |
| III | 2 | 22.39 | 22.30 | 22.48 | 22.38 |
| III | 3 | 4.13 | 1.26 | 2.06 | 0.74 |
| III | 4 | 1.11 | -0.36 | -0.99 | -0.76 |
| III | 5 | -2.33 | -16.62 | -19.39 | -17.88 |
| III | 6 | 19.58 | 9.22 | 10.70 | 5.57 |
| IV | 1 | 8.66 | 3.47 | 0.78 | 2.81 |
| IV | 2 | 4.66 | -8.69 | -14.18 | -12.63 |
| IV | 3 | 6.98 | -4.34 | -5.16 | -4.56 |
| IV | 4 | 7.97 | 8.40 | 7.49 | 8.33 |
| IV | 5 | 5.87 | -6.61 | -10.76 | -9.35 |
| IV | 6 | 15.84 | 15.42 | 14.63 | 15.32 |
| V | 1 | -2.22 | -13.84 | -13.72 | -13.73 |
| V | 2 | 15.95 | 16.13 | 16.09 | 16.43 |
| V | 3 | 1.04 | -3.93 | -4.55 | -4.08 |
| V | 4 | 25.22 | 25.27 | 25.28 | 24.96 |
| V | 5 | -10.66 | -12.74 | -12.43 | -12.25 |
| V | 6 | -1.81 | -14.05 | -17.56 | -16.25 |
| VI | 1 | 7.71 | -1.75 | -5.15 | -3.06 |
| VI | 2 | -1.79 | -9.24 | -10.08 | -9.47 |
| VI | 3 | 11.82 | 11.11 | 10.99 | 11.29 |
| VI | 4 | 10.34 | 8.12 | 6.84 | 7.51 |
| VI | 5 | 1.26 | -7.73 | -7.01 | -8.35 |
| VI | 6 | 8.24 | 5.42 | 4.24 | 5.56 |
| VII | 1 | 8.28 | -0.55 | -1.21 | -0.34 |
| VII | 2 | 2.32 | -1.29 | -2.35 | -1.55 |
| VII | 3 | 10.61 | 4.33 | 3.61 | 4.02 |
| VII | 4 | 7.58 | 3.21 | 3.09 | 2.95 |
| VII | 5 | 15.30 | 9.32 | 9.97 | 9.97 |
| VII | 6 | 4.05 | -1.81 | -4.67 | -2.72 |

#### Stan code

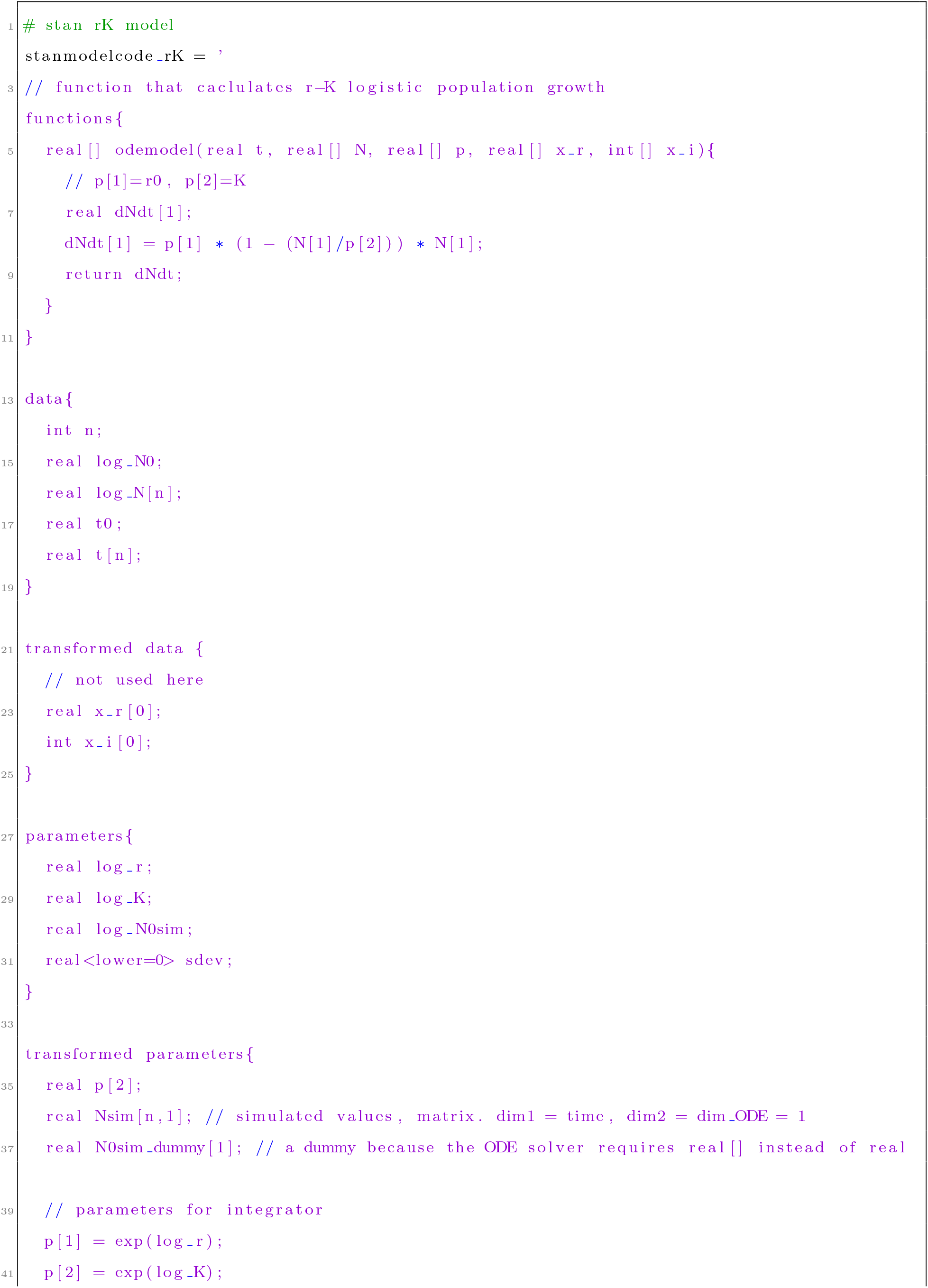
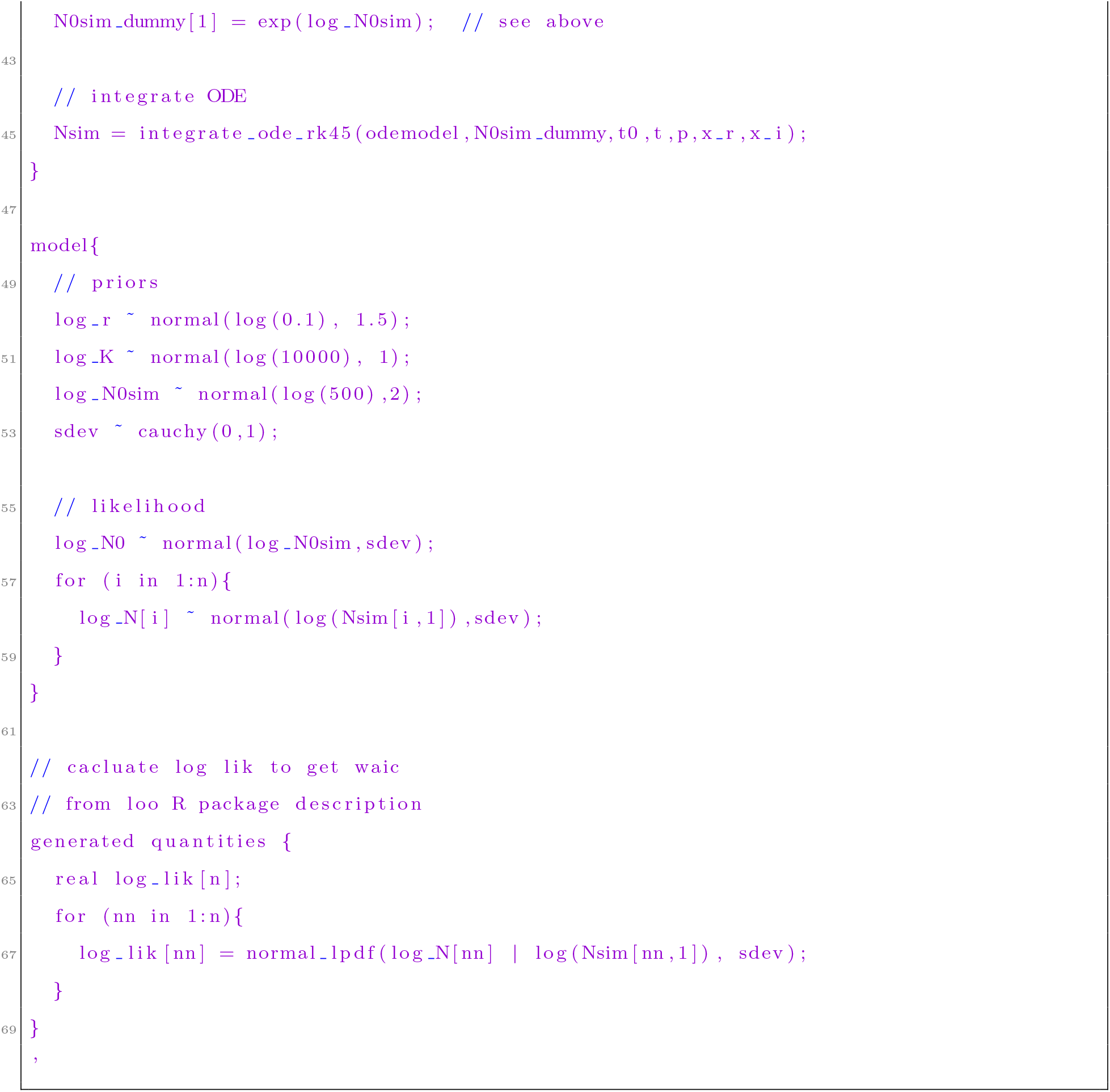
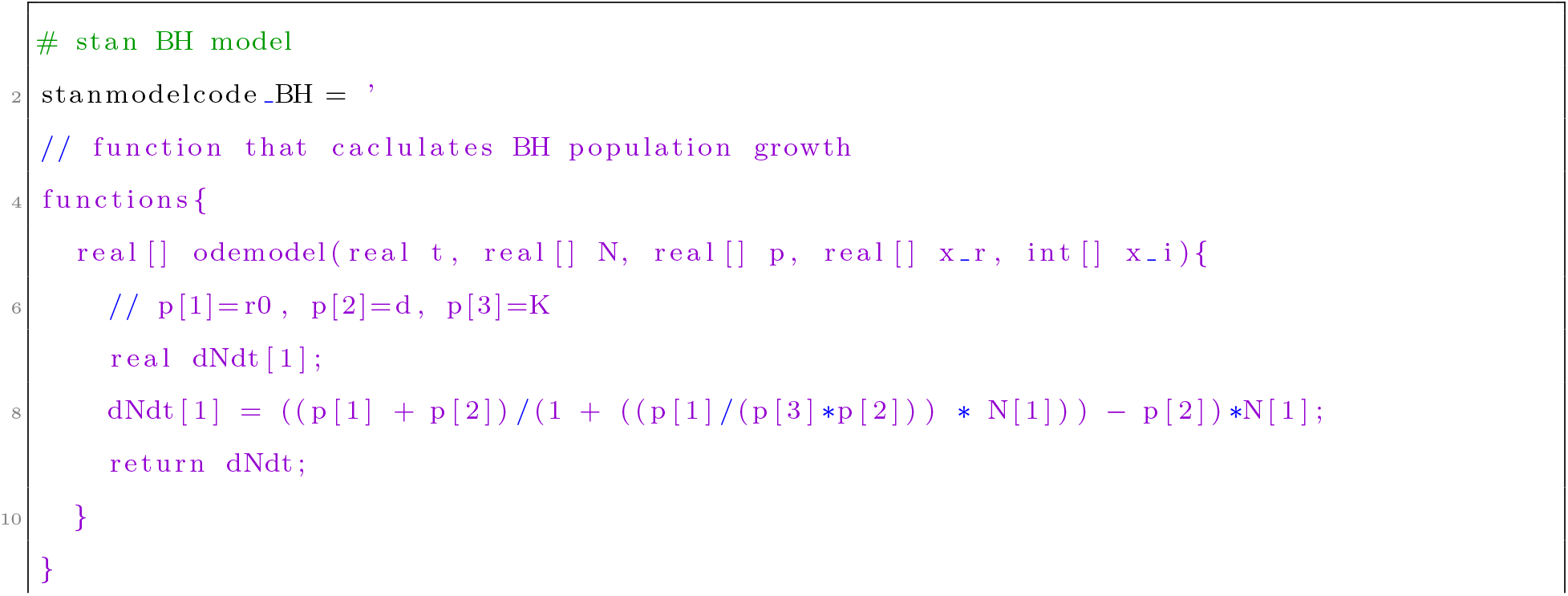
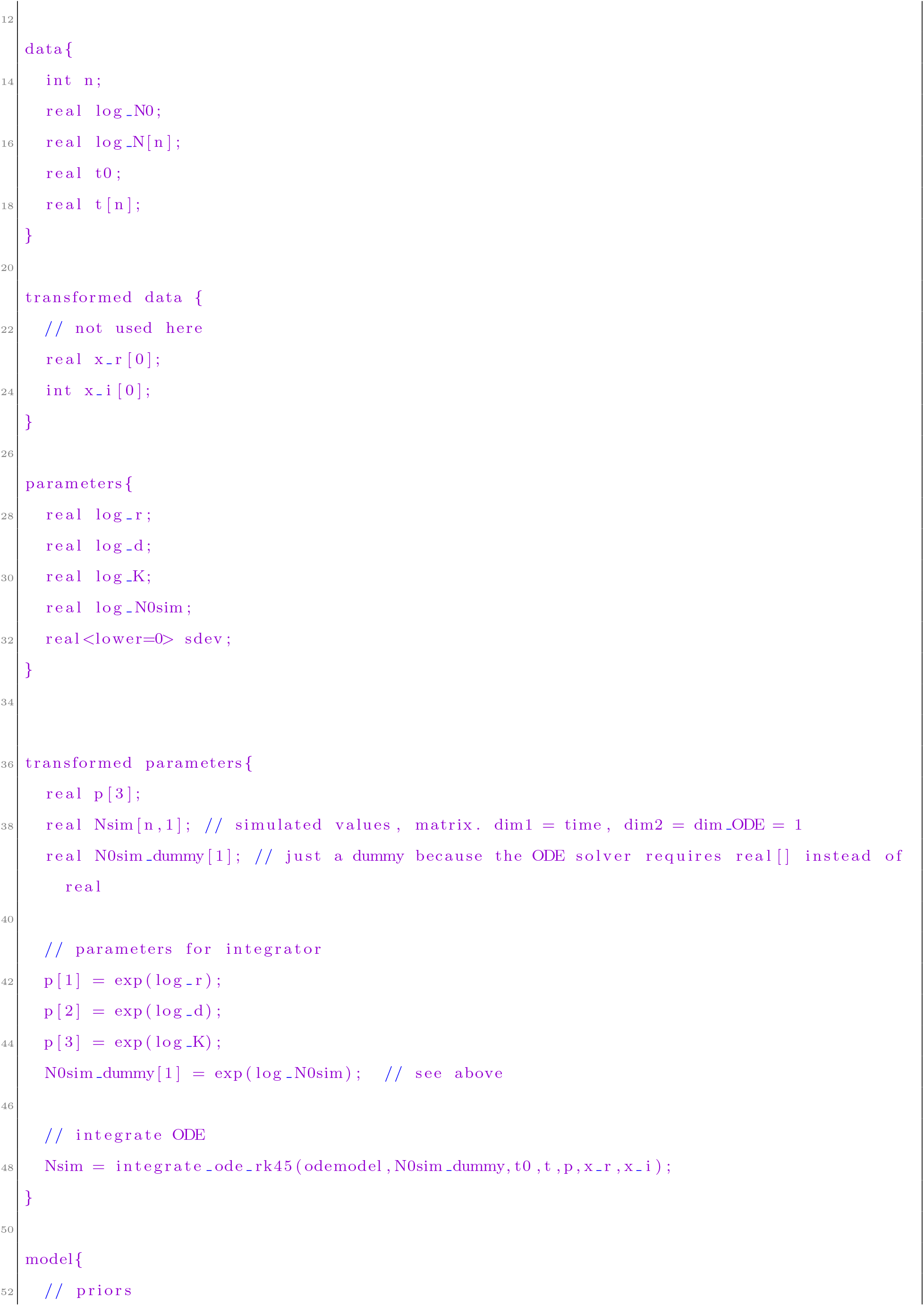
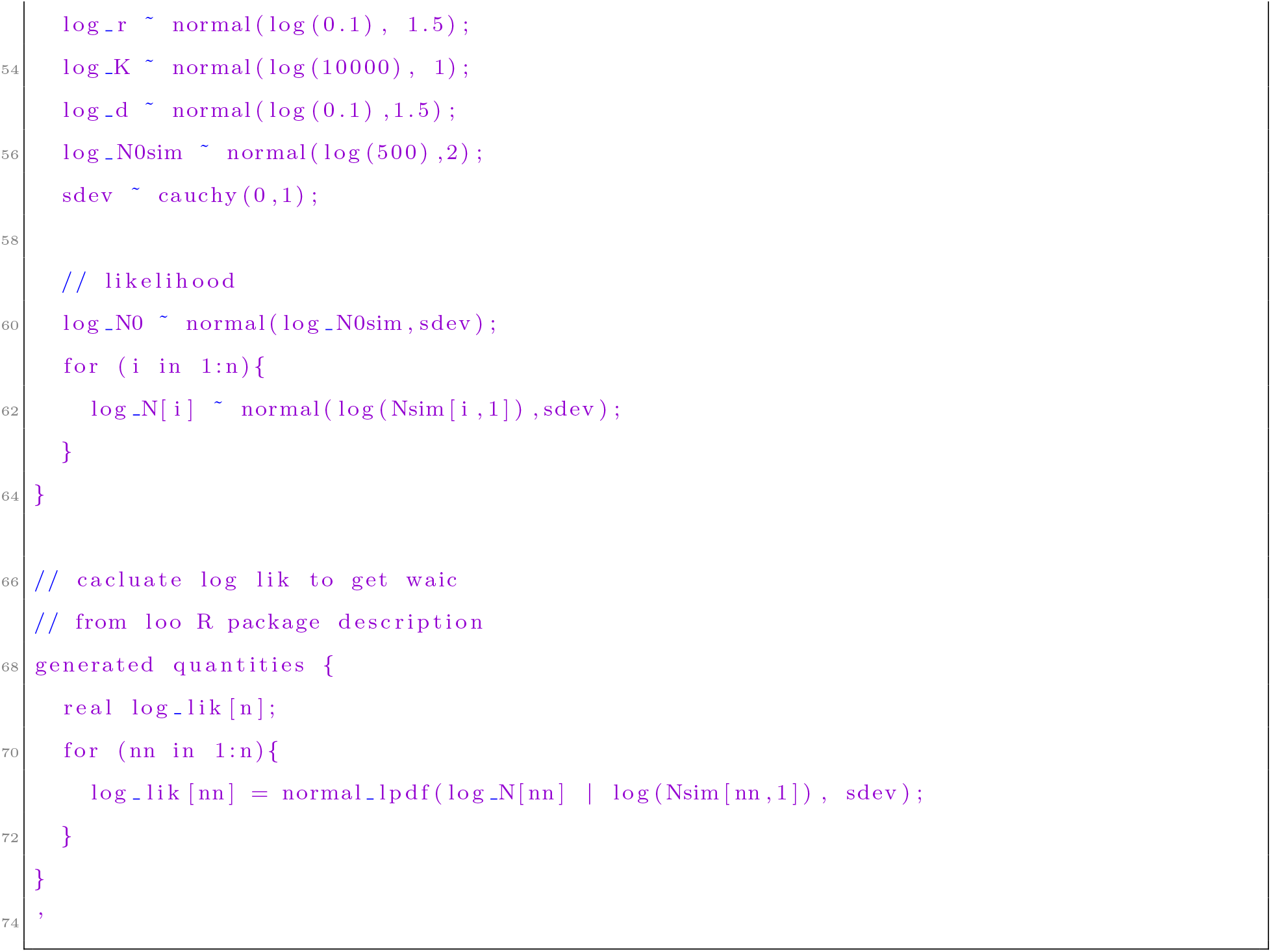
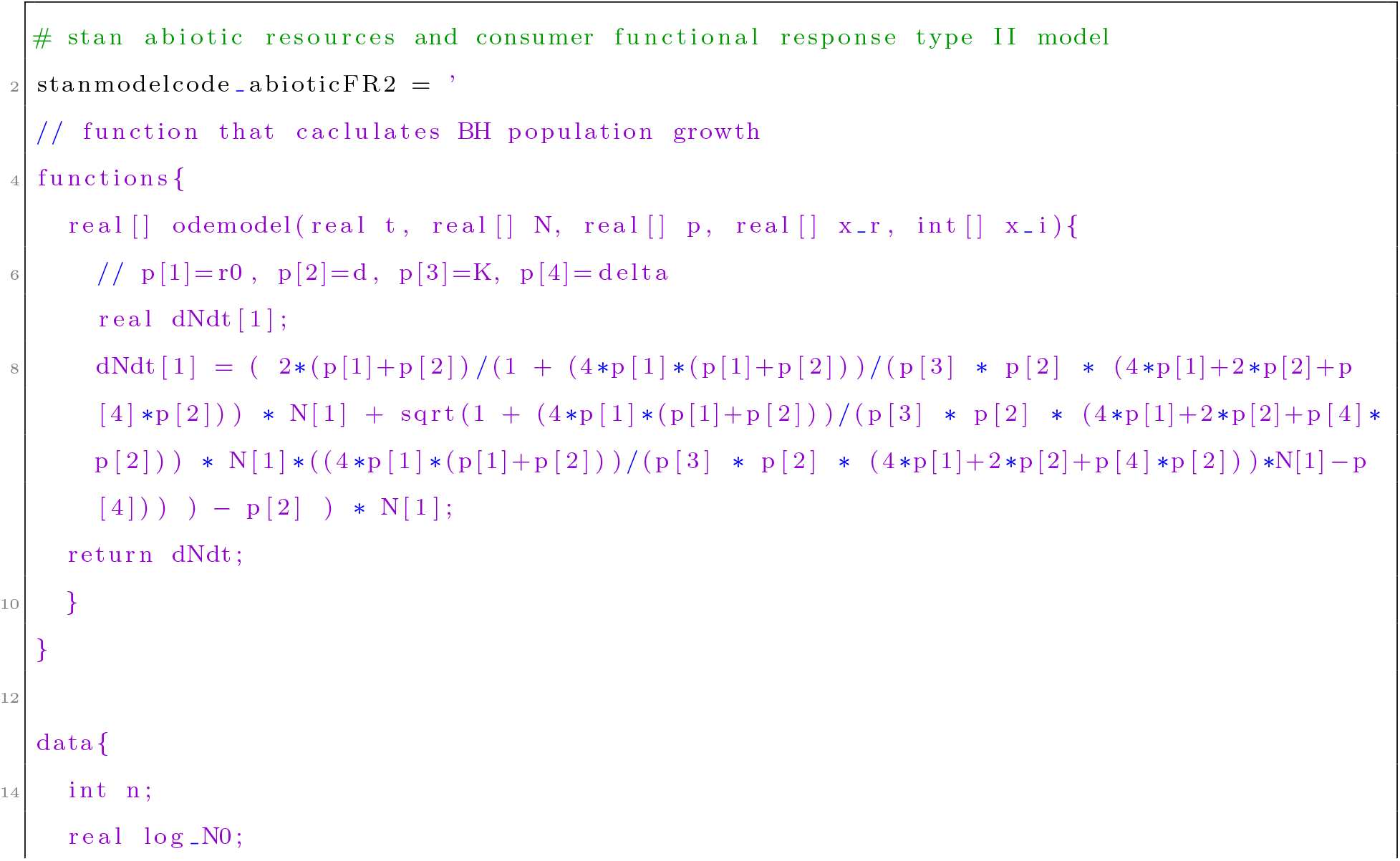
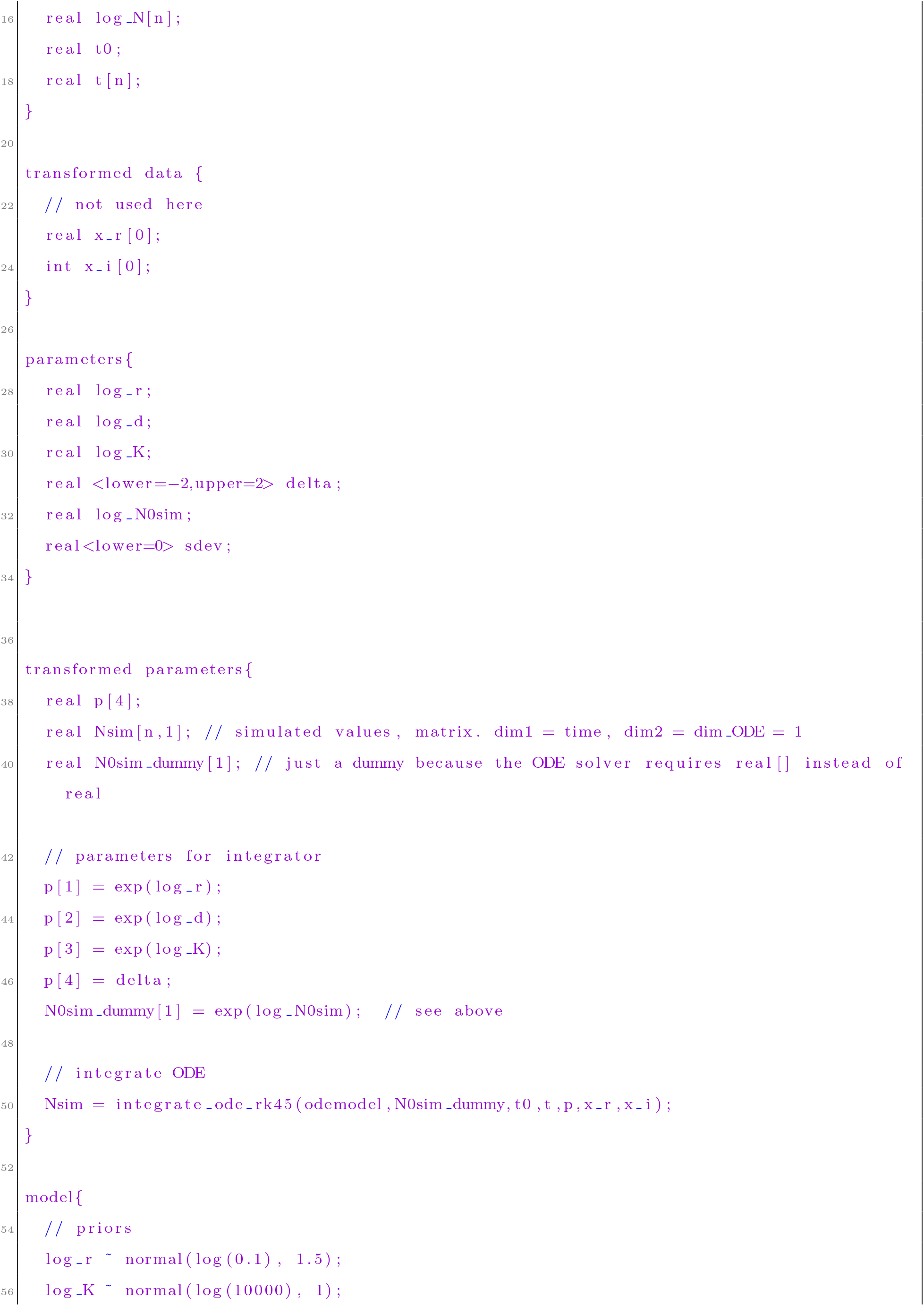
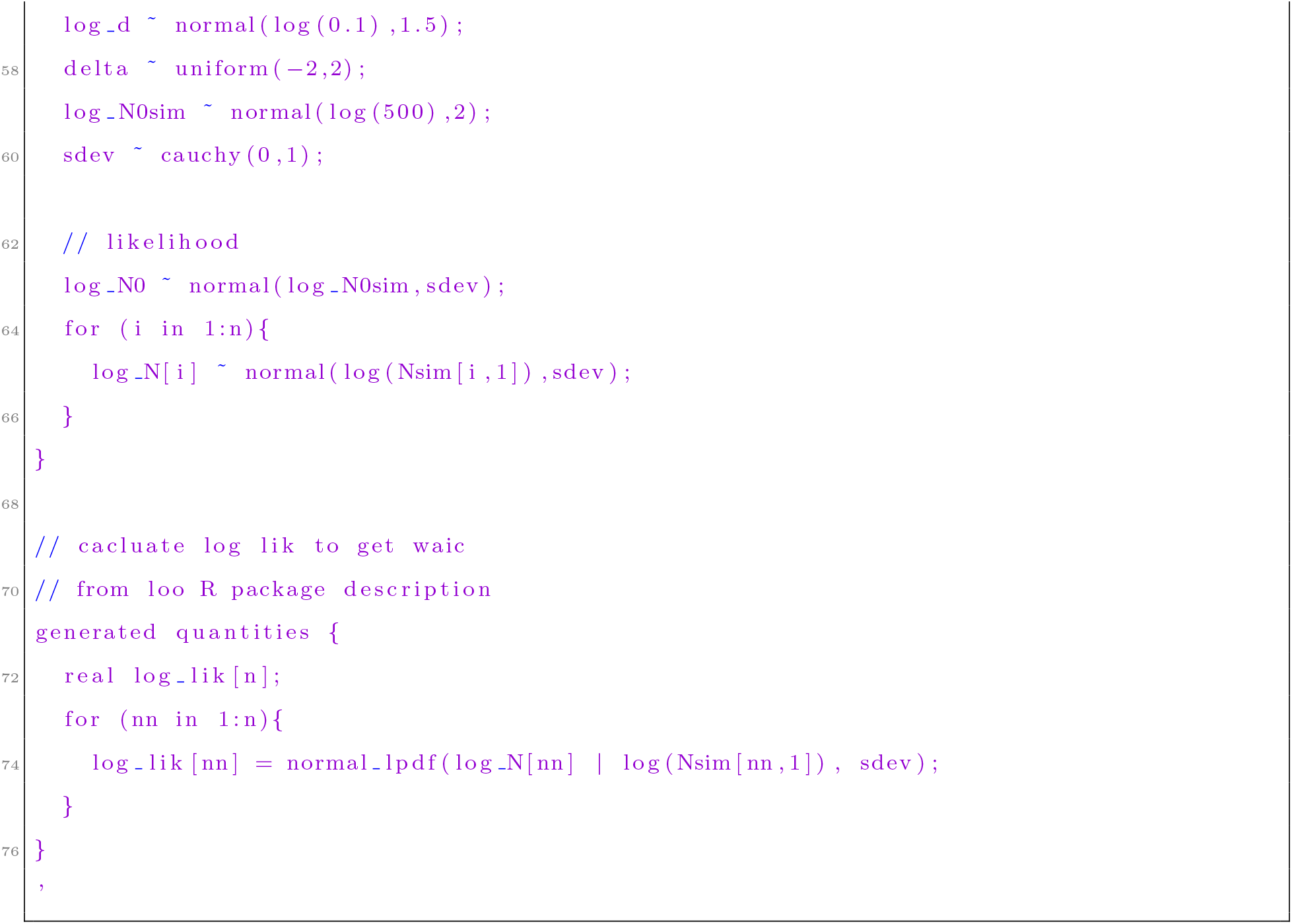
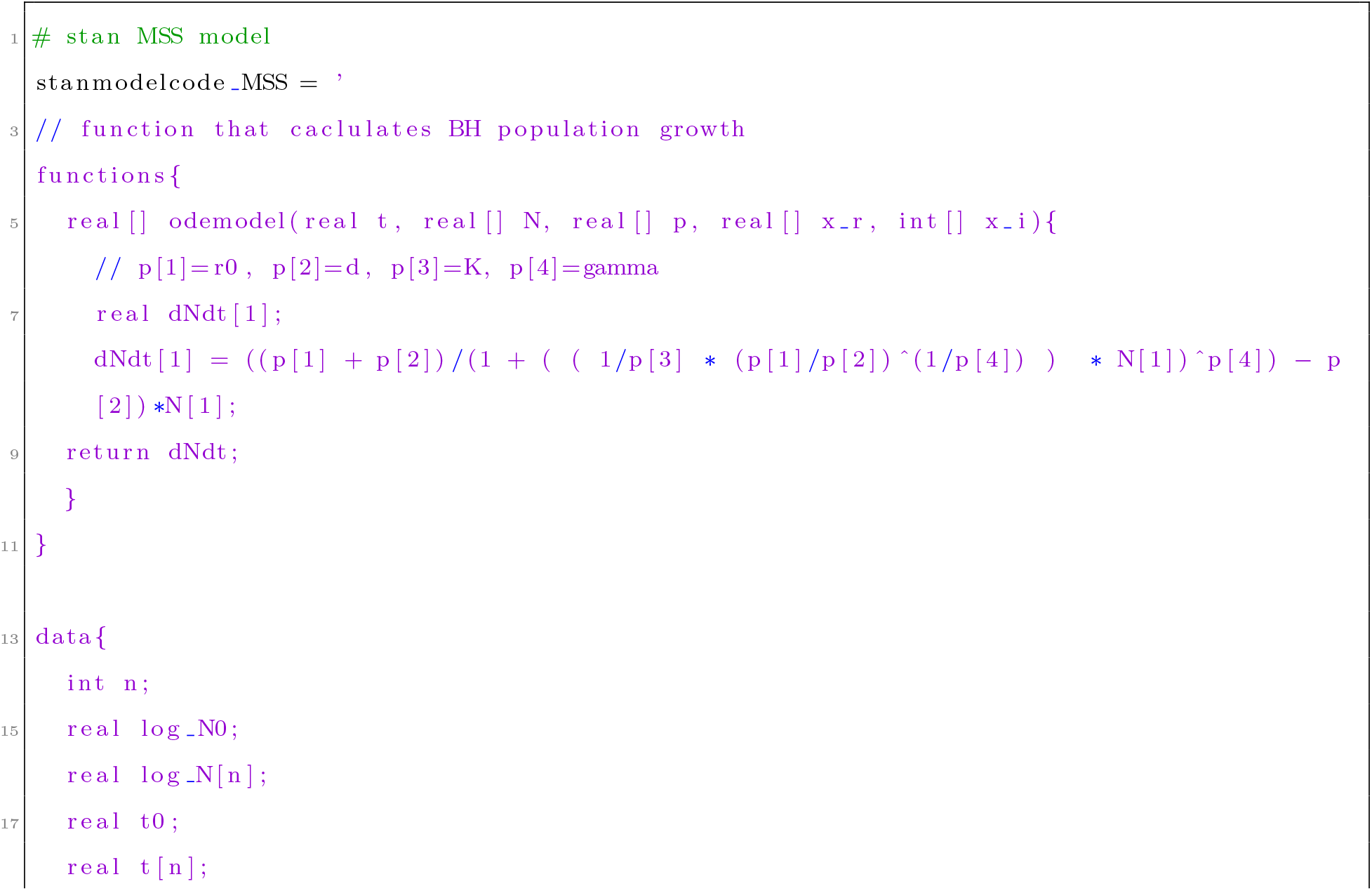
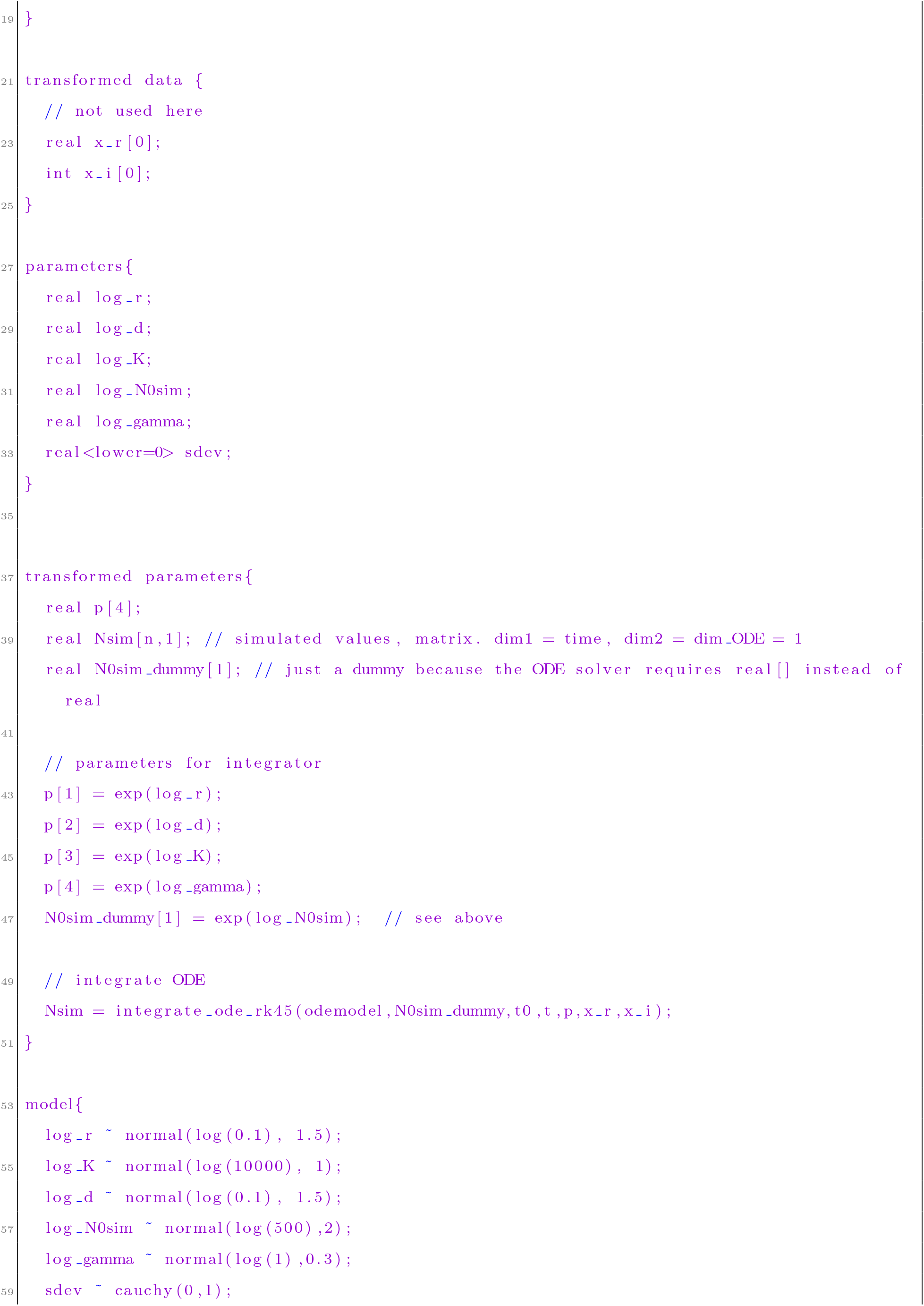
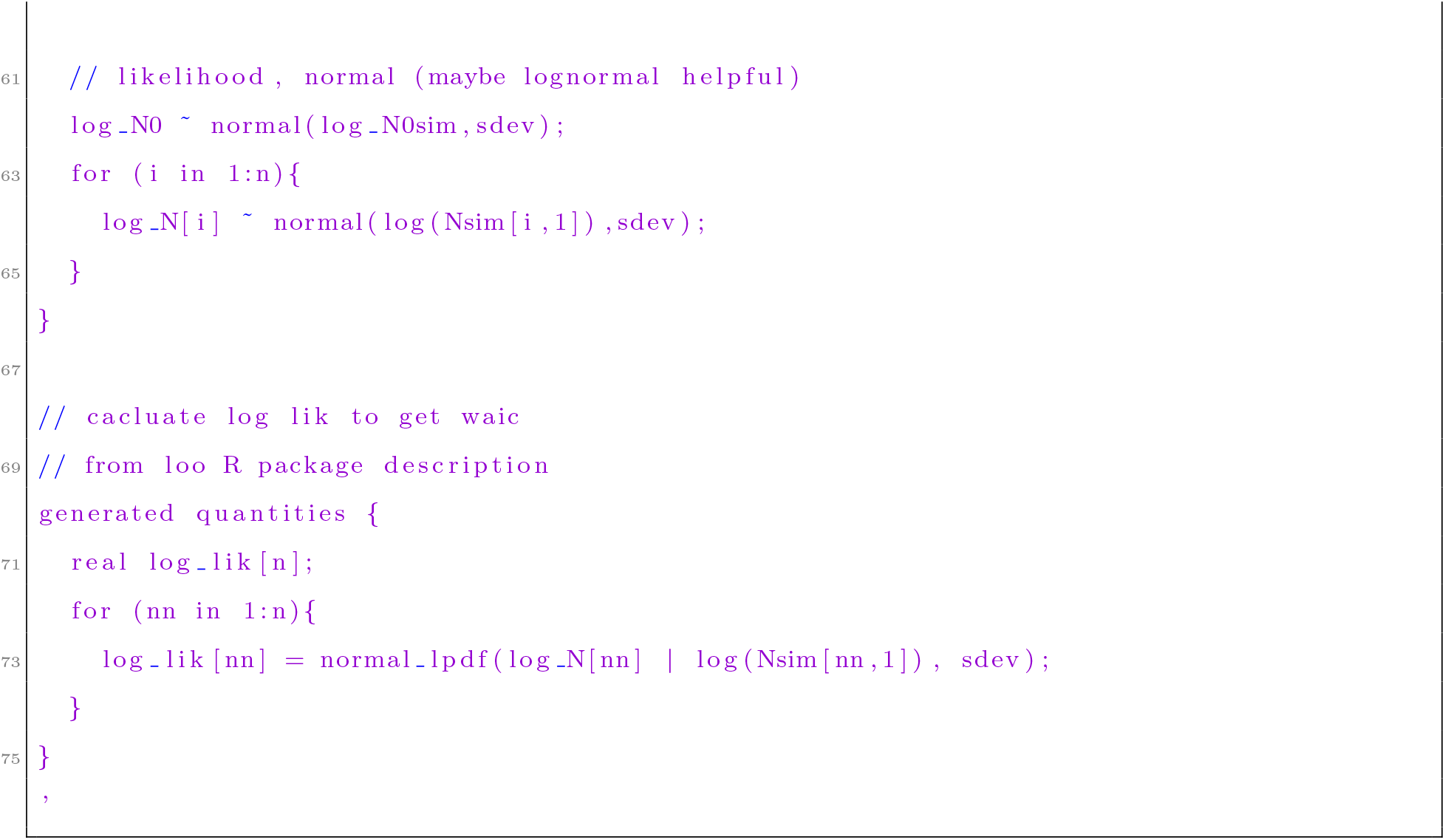

### S9 Temperature scaling of equilibrium densities

#### Overview

In order to analyse how the equilibrium densities change with temperature, we used metabolic theory of ecology (Brown et al., 2004) that states that a given biological rate *I* will depend on temperature as follows:

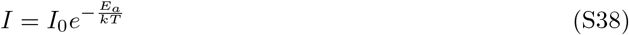

with *E_a_* as the activation energy, *k* the Boltzmann constant and *T* as the absolute temperature. While this relationship has been directly applied to population growth rates and competition coefficients, the effect of temperature on the equilibrium density has been less straight-forward to infer and justify theoretically (Gilbert et al., 2014; Bernhardt et al., 2018). Similarly to Uszko et al. (2017), our work provides a mechanistic way forward, at least in the case of abiotic resources. For non-saturating filter feeders, we can take the expression for equilibrium density in Eq. 10 and substitute Eq. S38 for all biological rates, that is, the feeding rate *a* and the death rate *d*, except for the efficiency *e*, which has been shown to be temperature independent (Del Giorgio and Cole, 1998; López-Urrutia and Morán, 2007). For simplicity, we assume the parameters of the abiotic resource to be temperature-independent and that all activation energies *E_a_* are identical. Then

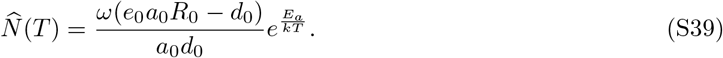

This result implies that, as temperature increases, equilibrium density decreases as shown in Fig. S8B for non-saturating filter feeding consumers feeding on abiotic resources (for details of the derivation see the Supplementary Material S9).

We can apply the same thoughts to the entire density-regulation function which then behaves as depicted in Fig. S8A. The simultaneous increase of population growth rate and competition coefficient with temperature decreases the equilibrium density and makes the density-regulation function steeper around the equilibrium density.

The negative scaling of equilibrium density with temperature holds more generally, for all cases of abiotic and biotic resources and shapes of functional responses investigated here (see Supplementary Material S9). Clearly, changing the temperature scaling law captured by Eq. S38 along the lines suggested by Uszko et al. (2017), that is, to be unimodal, because some rates may decline beyond certain critical temperature values, would change these results. While an in-depth analysis is beyond the scope of our current work, this extension highlights the potential of our framework. Of course, our derivations can be used to investigate the effects of other scaling laws, such as body size scaling, which we will not analyse any further here.

**Figure S8:**
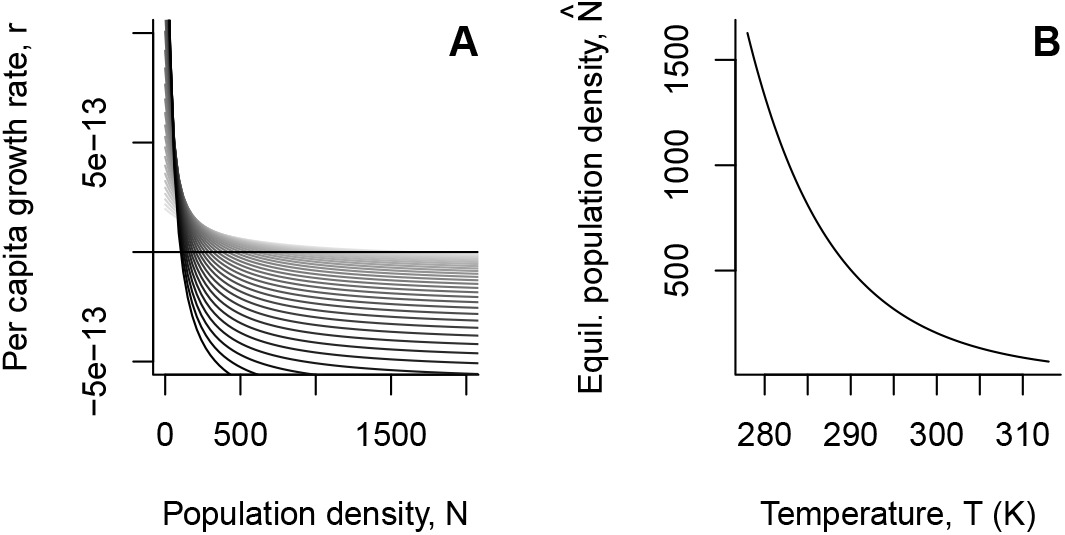
(A) Temperature scaling of the density-regulation function for non-saturating filter feeding consumers feeding on abiotic resources. With increasing temperature (darker grey) the density-regulation function becomes steeper because *r*_0_ and the competition coefficient *β* increase. (B) Temperature scaling of equilibrium population density. See the Supplementary Material S9 for a detailed derivation.

#### Detailed derivation

Substituting Eq. S38 for all biological rates, except the assimilation coefficient, e, which has been shown to be temperature independent (Del Giorgio and Cole, 1998; López-Urrutia and Morán, 2007), into Eq. 10 gives the expression of temperature dependent equilibrium population density of a non-saturating filter feeding consumer feeding on abiotic resource:

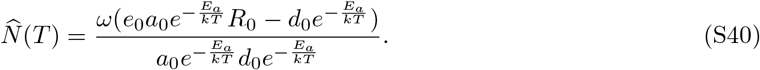

Assuming that all activation energies *E_a_* are identical, we can simplify this to

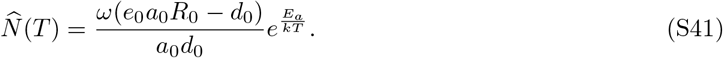

More generally, in terms of density dependence, we obtain for the capita growth rate of the consumer:

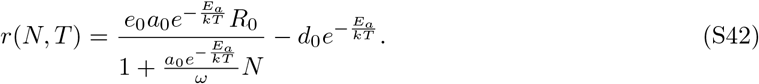

Under the same assumptions that all activation energies are identical, we can simplify the previous equation to:

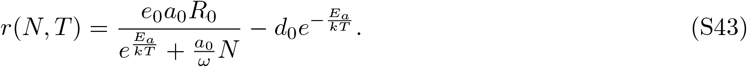

In analogy, temperature dependence of the equilibrium density can be investigated for type II functional responses and abiotic resources. In this case, we can use Eq. S11 and Eq. S38 to obtain:

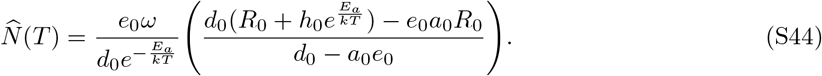

Similarly, we can use Eq. S32 and Eq. S38 to obtain the temperature dependence of a consumer characterized by a type II functional response feeding on biotic resources. Without repeating the above derivation we obtain 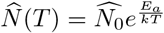.

### S10 Multi-species extensions

Our consideration below should be understood as representing examples and potential that can be developed. In general, developments should consider a systems logical consistency as detailed in Kuang (2002). We here again do not include predator-dependence of the functional responses (see Abrams, 2014, for a discussion). Further developments could also follow ideas laid out by Bastolla et al. (2005).

#### Overview

Considering our simplest case, abiotic resources and a non-saturating filter feeding consumer, we can add *n* consumer species to the system (see Ruggieri and Schreiber, 2005, for an example with two consumers). After solving for resource equilibrium equilibrium, substitution and some reparametrization (see Supplementary Material S10), we obtain the consumer dynamics for species *i*:

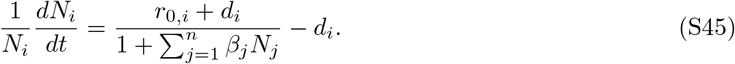

Based on Eq. S45, coexistence is only possible if growth and death rates are identical, because all consumers feed on one resource and do not interact in any other way. In analogy to Matessi and Gatto (1984) the species that minimizes 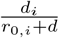 excludes the others (see also Hofbauer and Sigmund, 1998).

If there is any interspecific interaction that is not resource-mediated, an additional interaction term may be added in analogy to a death term as follows:

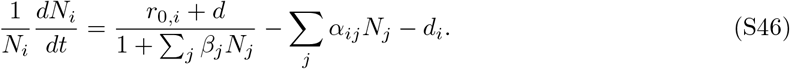

This additional density-dependent mortality term allows for coexistence of more than one species (McPeek, 2012). Possible mechanisms behind the additional interaction term include cannibalism at the intraspecific level and chemical compounds (that is, allelopathy) at both the intra- and interspecific levels.

Finally, for more than one (abiotic) resource and more than one (non-saturating filter feeding) consumer we obtain for species i after simplification:

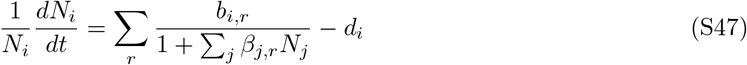

where *b_i,r_* = *e_i,r_ a_i,r_ R*_0,*r*_ is the resource specific birth term of species *i* and 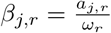 is the competition coefficient. See below for details.

#### Multiple consumers

We here consider consumer-resource dynamics, where we have two consumer species feeding on one abiotic resource (similar to Eqs. 5 and 6 in the main text). In analogy to our previous considerations, we assume the following dynamics:

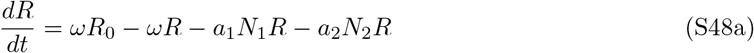

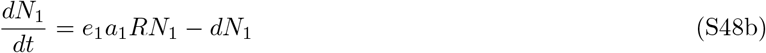

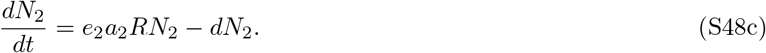

Solving the resource for equilibrium we get:

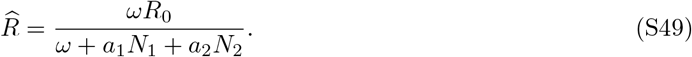

We can now substitute Eq. S49 into Eq. S48b and get:

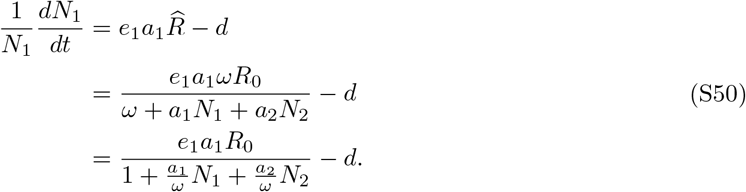

Similarly, substituting equation (S49) into equation (S48c) gives:

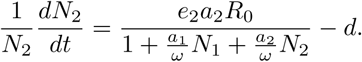

Hence, for a system with *n* consumer species, uniquely indexed with *i* ∈ {1, …, *n*}, we get the consumer dynamics of species *i*:

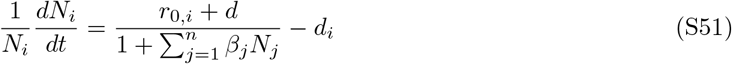

with 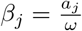.

#### Multiple resources

There may also exist consumer-resource systems with multiple resources. We here consider a system with two abiotic resources, and one consumer species. This results in the following system of equations:

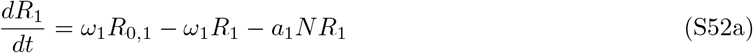

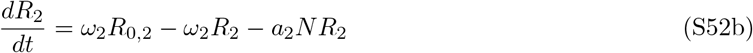

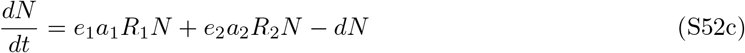

Solving this system to resource equilibrium gives for 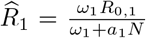 and 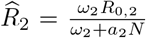. Substituting both resource equilibria into Eq. S52C gives:

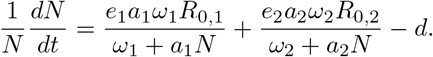

Hence, for a system with multiple resources, uniquely indexed with *r* ∈ {1, …, *m*}, we would find the following consumer dynamics:

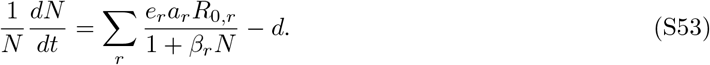

However, when determining *r*_0_ by setting *N* = 0, we get: 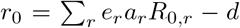. Note that because of the different denominators we cannot easily substitute this into Eq. S53.

#### Multiple resources and multiple consumers

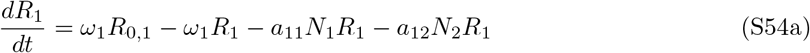

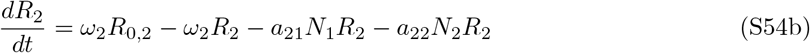

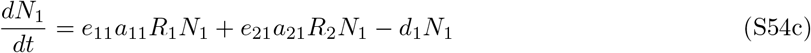

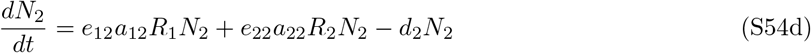

Solving to resource equilibrium gives 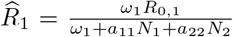 and 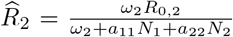 Substituting these resource equilibria into Eq. S54c gives:

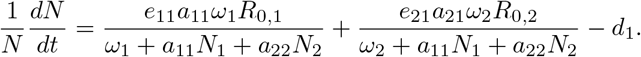

Similarly, substituting 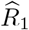 and 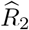 into Eq. S54d gives:

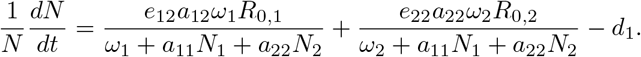

From this it follows that for a system with *m* resources and *n* consumers, the consumer dynamics of a consumer species *i* is given by the following equation:

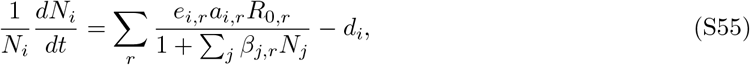

where 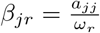. This is Eq. S47 found in the main text.

## References

Abrams, P. A. 1975. Limiting similarity and the form of the competition coefficient. – Theor. Popul. Biol. 8: 356–375.

Abrams, P. A. 1977. Density-Independent Mortality and Interspecific Competition: A Test of Pianka’s Niche Overlap Hypothesis. – Am. Nat. 111(979): 539–552.

Abrams, P. A. 1988. Resource Productivity-Consumer Species Diversity: Simple Models of Competition in Spatially Heterogeneous Environments. – Ecology 69(5): 1418–1433.

Abrams, P. A. 2002. Will Small Population Sizes Warn Us of Impending Extinctions?. – Am. Nat. 160(3): 293–305.

Abrams, P. A. 2009a. Adaptive changes in prey vulnerability shape the response of predator populations to mortality. – J. Theor. Biol. 261(2): 294–304.

Abrams, P. A. 2009b. Determining the Functional Form of Density Dependence: Deductive Approaches for Consumer Resource Systems Having a Single Resource. – Am. Nat. 174(3): 321–330.

Abrams, P. A. 2009c. The implications of using multiple resources for consumer density dependence. – Evol. Ecol. Res. 11(4): 517–540.

Abrams, P. A. 2014. Why ratio dependence is (still) a bad model of predation. – Biol. Rev. 90(3): 794–814.

Abrams, P. A. 2019. How Does the Evolution of Universal Ecological Traits Affect Population Size? Lessons from Simple Models. – Am. Nat. 193(6): 814–829.

Aktipis, C. A., Boddy, A. M., Gatenby, R. A., Brown, J. S. and Maley, C. C. 2013. Life history trade-offs in cancer evolution. – Nat. Rev. Cancer 13(12): 883.

Allee, W. C. 1931. Animal aggregations: a study in general sociology. – University of Chicago Press.

Begon, M., Townsend, C. R. and Harper, J. L. 2006. Ecology: From Individuals to Ecosystems. – Blackwell Publishing.

Bellows, T. S. 1981. The descriptive properties of some models for density dependence. – J. Anim. Ecol. 50: 139–156.

Bernhardt, J. R., Sunday, J. M. and O’Connor, M. I. 2018. Metabolic theory and the temperature-size rule explain the temperature dependence of population carrying capacity. – Am. Nat. 192(6): 687–697.

Beverton, R. J. H. and Holt, S. J. 1957. On the dynamics of exploited fish populations. – Chapman & Hall.

Borlestean, A., Frost, P. C. and Murray, D. L. 2015. A mechanistic analysis of density dependence in algal population dynamics. – Front. Ecol. Evol. 3.

Brännström, A. and Sumpter, D. J. T. 2005. The role of competition and clustering in population dynamics. – Proc. R. Soc. B-Biol. Sci. 272(1576): 2065–2072.

Brown, J. H., Gillooly, J. F., Allen, A. P., Savage, V. M. and West, G. B. 2004. Toward a metabolic theory of ecology. – Ecology 85(7): 1771–1789.

Burton, O. J., Pillips, B. L. and Travis, J. M. J. 2010. Trade-offs and the evolution of life-histories during range expansion. – Ecol. Lett. 13(10): 1210–1220.

Clark, F., Brook, B. W., Delean, S., Reit Akakaya, H. and Bradshaw, C. J. A. 2010. The theta-logistic is unreliable for modelling most census data. – Methods Ecol. Evol. 1(3): 253–262.

Cosner, C., DeAngelis, D. L., Ault, J. S. and Olson, D. B. 1999. Effects of spatial grouping on the functional response of predators. – Theor. Popul. Biol. 56(1): 65–75.

Courchamp, F., Berec, L. and Gascoigne, J. 2008. Allee Effects in Ecology and Conservation. – Oxford University Press.

Dallas, T., Decker, R. R. and Hastings, A. 2017. Species are not most abundant in the centre of their geographic range or climatic niche. – Ecol. Lett. 20(12): 1526–1533.

Delean, S., Brook, B. W. and Bradshaw, C. J. A. 2012. Ecologically realistic estimates of maximum population growth using informed Bayesian priors. – Methods Ecol. Evol. 4(1): 34–44.

Eberhardt, L. L., Breiwick, J. M. and Demaster, D. P. 2008. Analyzing population growth curves. – Oikos 117: 1240–1246.

Engen, S. and Sæther, B.-E. 2017. r-and K-selection in fluctuating populations is determined by the evolutionary trade-off between two fitness measures: Growth rate and lifetime reproductive success. – Evolution 71(1): 167–173.

Fleischer, S. R., Li, J. et al. 2018. Pick your trade-offs wisely: Predator-prey eco-evo dynamics are qualitatively different under different trade-offs. – J. Theor. Biol. 456: 201–212.

Fronhofer, E. A. and Altermatt, F. 2015. Eco-evolutionary feedbacks during experimental range expansions. – Nat. Commun. 6: 6844.

Fronhofer, E. A., Nitsche, N. and Altermatt, F. 2017. Information use shapes the dynamics of range expansions into environmental gradients. – Glob. Ecol. Biogeogr. 26(4): 400–411.

Gabriel, J.-P., Saucy, F. and Bersier, L.-F. 2005. Paradoxes in the logistic equation?. – Ecol. Model. 185(1): 147–151.

Gatto, M. 1990. General Minimum Principle for Competing Populations: Some Ecological and Evolutionary Consequences. – Theor. Popul. Biol. 37: 369–388.

Geritz, S. and Gyllenberg, M. 2012. A mechanistic derivation of the DeAngelisBeddington functional response. – J. Theor. Biol. 314(7): 106–108.

Geritz, S. A. H. and Kisdi, E. 2004. On the mechanistic underpinning of discrete-time population models with complex dynamics. – J. Theor. Biol. 228(2): 261–269.

Getz, W. 1993. Metaphysiological and evolutionary dynamics of populations exploiting constant and interactive resources:RK selection revisited. – Evol. Ecol. 7(3): 287–305.

Ghedini, G., Loreau, M., White, C. R. and Marshall, D. J. 2018. Testing MacArthur’s minimisation principle: do communities minimise energy wastage during succession?. – Ecol. Lett. 21: 1182–1190.

Gilbert, B., Tunney, T., McCann, K., DeLong, J., Vasseur, D., Savage, V., Shurin, J. B., Dell, A. I., Barton, B. T., Harley, C. D. G., Kharouba, H. M., Kratina, P., Blanchard, J. L., Clements, C., Winder, M., Greig, H. S. and O’Connor, M. I. 2014. A bioenergetic framework for the temperature dependence of trophic interactions. – Ecol. Lett. 17: 902–914.

Gilpin, M. E. and Ayala, F. J. 1973. Global Models of Growth and Competition. – Proc. Natl. Acad. Sci. U. S. A. 70(12): 3590–3593.

Ginzburg, L. R. 1992. Evolutionary consequences of basic growth equations. – Trends Ecol. Evol. 7(4): 133.

Govaert, L., Fronhofer, E. A., Lion, S., Eizaguirre, C., Bonte, D., Egas, M., Hendry, A. P., Martins, A. D. B., Melia’n, C. J., Raeymaekers, J., Ratikainen, I. I., Saether, B.-E., Schweitzer, J. A. and Matthews, B. 2019. Eco-evolutionary feedbacks – theoretical models and perspectives. – Funct. Ecol. 33(1): 13–30.

Hendry, A. P. 2017. Eco-evolutionary dynamics. – Princeton University Press.

Henle, K., Sarre, S. and Wiegand, K. 2004. The role of density regulation in extinction processes and population viability analysis. – Biodivers. Conserv. 13(1): 9–52.

Herrando-Prez, S., Delean, S., Brook, B. W. and Bradshaw, C. J. A. 2012. Density dependence: an ecological Tower of Babel. – Oecologia 170(3): 585–603.

Hiltunen, T., Hairston, N. G., Hooker, G., Jones, L. E. and Ellner, S. P. 2014. A newly discovered role of evolution in previously published consumer-resource dynamics. – Ecol. Lett. 17(8): 915–923.

Hodgson, E. E., Essington, T. E. and Halpern, B. S. 2017. Density dependence governs when population responses to multiple stressors are magnified or mitigated. – Ecology 98(10): 2673–2683.

Holling, C. S. 1959. The components of predation as revealed by a study of small-mammal predation of the European pine sawfly. – Can. Entomol. 91(05): 293–320.

Jeschke, J. M., Kopp, M. and Tollrian, R. 2004. Consumer-food systems: why type I functional responses are exclusive to filter feeders. – Biol. Rev. 79(2): 337–349.

Johst, K., Berryman, A. and Lima, M. 2008. From individual interactions to population dynamics: individual resource partitioning simulation exposes the causes of nonlinear intra-specific competition. – Popul. Ecol. 50(1): 79–90.

Joshi, A. and Mueller, L. D. 1988. Evolution of higher feeding rate in Drosophila due to density-dependent natural selection. – Evolution 42(5): 1090–1093.

Joshi, A. and Mueller, L. D. 1996. Density-dependent natural selection in Drosophila: Trade-offs between larval food acquisition and utilization. – Evol. Ecol. 10(5): 463–474.

Joshi, A., Prasad, N. G. and Shakarad, M. 2001. K-selection, α-selection, effectiveness, and tolerance in competition: Density-dependent selection revisited. – J. Genet. 80(2): 63–75.

Kostitzin, V. A. 1937. Biologie mathématique. – Armand Colin.

Krebs, C. J. 2015. One hundred years of population ecology: successes, failures, and the road ahead. – Integr. Zool. 10: 233–240.

Kubisch, A., Winter, A.-M. and Fronhofer, E. A. 2016. The downward spiral: eco-evolutionary feedback loops lead to the emergence of ‘elastic’ ranges. – Ecography 39(3): 261–269.

Kurtz, T. G. 1981. Approximation of population processes. – vol. 36. SIAM.

Lakin, W. and Van Den Driessche, P. 1977. Time Scales in Population Biology. – SIAM J. Appl. Math. 32(3): 694–705.

Lande, R., Engen, S. and Saether, B. E. 2009. An evolutionary maximum principle for density-dependent population dynamics in a fluctuating environment. – Philos. Trans. R. Soc. B-Biol. Sci. 364(1523): 1511–1518.

Luckinbill, L. S. 1979. Selection and the r/K continuum in experimental populations of protozoa. – Am. Nat. 113(3): 427–437.

MacArthur, R. 1969. Species packing, and what interspecies competition minimizes. – Proc. Natl. Acad. Sci. U. S. A. 64(4): 1396–1371.

MacArthur, R. 1970. Species packing and competitive equilibrium for many species. – Theor. Popul. Biol. 1(1): 1–11.

MacArthur, R. and Levins, R. 1967. The Limiting Similarity, Convergence, and Divergence of Coexisting Species. – Am. Nat. 101(921): 377–385.

MacArthur, R. H. 1962. Some generalized theorems of natural selection. – Proc. Natl. Acad. Sci. U. S. A. 48(11): 1893–1897.

Mallet, J. 2012. The struggle for existence: how the notion of carrying capacity, K, obscures the links between demography, Darwinian evolution, and speciation. – Evol. Ecol. Res. 14(5): 627–665.

Matessi, C. and Gatto, M. 1984. Does K-selection imply prudent predation?. – Theor. Popul. Biol. 25(3): 347–363.

Maynard Smith, J. and Slatkin, M. 1973. The Stability of Predator-Prey Systems. – Ecology 54(2): 384–391.

McElreath, R. 2016. Statistical Rethinking: a Bayesian course with examples in R and Stan. – Chapman & Hall/CRC.

McPeek, M. A. 2017. The Ecological Dynamics of Natural Selection: Traits and the Coevolution of Community Structure. – Am. Nat. 189(5): E91–E117.

Melbourne, B. A. and Hastings, A. 2009. Highly variable spread rates in replicated biological invasions: fundamental limits to predictability. – Science 325(5947): 1536–1539.

Mobilia, M., Georgiev, I. T. and Täuber, U. C. 2007. Phase transitions and spatio-temporal fluctuations in stochastic lattice Lotka-Volterra models. – J. Stat. Phys. 128(1-2): 447–483.

Mueller, L., Guo, P. and Ayala, F. 1991. Density-dependent natural selection and trade-offs in life history traits. – Science 253(5018): 433–435.

Mueller, L. D. 1990. Density-dependent natural selection does not increase efficiency. – Evol. Ecol. 4(4): 290–297.

Mueller, L. D. 1997. Theoretical and empirical examination of density-dependent selection. – Annu. Rev. Ecol. Syst. pp. 269–288.

O’Dwyer, J. P. 2018. Whence Lotka-Volterra?. – Theor. Ecol. 11(4): 441–452.

Palkovacs, E. P., Wasserman, B. A. and Kinnison, M. T. 2011. Eco-Evolutionary Trophic Dynamics: Loss of Top Predators Drives Trophic Evolution and Ecology of Prey. – PLoS ONE 6(4): e18879.

Pástor, L., Botta-Dukát, Z., Magyar, G., Czárán, T. and Meszéna, G. 2016. Theory-based ecology: A Darwinian approach. – Oxford University Press.

Ratikainen, I. I., Gill, J. A., Gunnarsson, T. G., Sutherland, W. J. and Kokko, H. 2008. When density dependence is not instantaneous: theoretical developments and management implications. – Ecol. Lett. 11(4): 184–198.

Reding-Roman, C., Hewlett, M., Duxbury, S., Gori, F., Gudelj, I. and Beardmore, R. 2017. The unconstrained evolution of fast and efficient antibiotic-resistant bacterial genomes. – Nat. Ecol. Evol. 1(3): 0050.

Reynolds, S. A. and Brassil, C. E. 2013. When can a single-species, density-dependent model capture the dynamics of a consumer-resource system?. – J. Theor. Biol. 339: 70–83.

Reznick, D., Bryant, M. J. and Bashey, F. 2002. r-and K-selection revisited: the role of population regulation in life-history evolution. – Ecology 83(6): 1509–1520.

Reznick, D. and King, K. 2017. Antibiotic resistance: Evolution without trade-offs. – Nat. Ecol. Evol. 1(3): 0066.

Rosenbaum, B., Raatz, M., Weithoff, G., Fussmann, G. F. and Gaedke, U. 2019. Estimating parameters from multiple time series of population dynamics using Bayesian inference. – Front. Ecol. Evol. 6: 234.

Ross, J. 2009. A note on density dependence in population models. – Ecol. Model. 220(23): 3472–3474.

Rueffler, C., Egas, M. and Metz, J. A. J. 2006. Evolutionary Predictions Should Be Based on IndividualLevel Traits. – Am. Nat. 168(5): E148–E162.

Ruggieri, E. and Schreiber, S. J. 2005. The dynamics of the Schoener-Polis-Holt model of intra-guild predation. – Math. Biosci. Eng. 2(2): 279–288.

Saether, B.-E., Visser, M. E., Grtan, V. and Engen, S. 2016. Evidence for r-and K-selection in a wild bird population: a reciprocal link between ecology and evolution. – Proc. R. Soc. B-Biol. Sci. 283(1829): 20152411.

Schoener, T. W. 1973. Population growth regulated by intraspecific competition for energy or time: Some simple representations. – Theor. Popul. Biol. 4(1): 56–84.

Schoener, T. W. 1974. Some Methods for Calculating Competition Coefficients from Resource-Utilization Spectra. – Am. Nat. 108(961): 332–340.

Schoener, T. W. 1978. Effects of density-restricted food encounter on some single-level competition models. – Theor. Popul. Biol. 13(3): 365–381.

Sibly, R. M., Barker, D., Denham, M. C., Hone, J. and Pagel, M. 2005. On the Regulation of Populations of Mammals, Birds, Fish, and Insects. – Science 309(5734): 607–610.

Thieme, H. R. 2003. Mathematics in Population Biology. – Princeton University Press.

Tilman, D. 1980. Resources: A Graphical-Mechanistic Approach to Competition and Predation. – Am. Nat. 116(3): 362–393.

Travis, J. M. J., Delgado, M., Bocedi, G., Baguette, M., Barto, K., Bonte, D., Boulangeat, I., Hodgson, J. A., Kubisch, A., Penteriani, V., Saastamoinen, M., Stevens, V. M. and Bullock, J. M. 2013. Dispersal and species responses to climate change. – Oikos 122(11): 1532–1540.

Turchin, P. 1999. Population Regulation: A Synthetic View. – Oikos 84(1): 153.

Turchin, P. 2003. Complex Population Dynamics: A Theoretical/Empirical Synthesis. – Princeton University Press.

Uszko, W., Diehl, S., Englund, G. and Amarasekare, P. 2017. Effects of warming on predator-prey interactions – a resource-based approach and a theoretical synthesis. – Ecol. Lett. 20(4): 513–523.

Verhulst, P.-F. 1838. Notice sur la loi que la population suit dans son accroissement. – Correspondance Math’ematique et Physique 10: 113–121.

Wei, X. and Zhang, J. 2019. Environment-dependent pleiotropic effects of mutations on the maximum growth rate r and carrying capacity K of population growth. – PLoS Biol. 17(1): e3000121.

Yodzis, P. and Innes, S. 1992. Body size and consumer-resource dynamics. – Am. Nat. 139(6): 1151–1175.

Yoshida, T., Jones, L. E., Ellner, S. P., Fussmann, G. F. and Hairston, N. G. 2003. Rapid evolution drives ecological dynamics in a predatorprey system. – Nature 424(6946): 303–306.

## Supplementary References

Abrams, P. A. 2009. Determining the Functional Form of Density Dependence: Deductive Approaches for ConsumerResource Systems Having a Single Resource. – Am. Nat. 174(3): 321–330.

Altermatt, F. and Fronhofer, E. A. 2018. Dispersal in dendritic networks: ecological consequences on the spatial distribution of population densities. – Freshwater Biol. 63(1): 22–32.

Bastolla, U., Lässig, M., Manrubia, S. C. and Valleriani, A. 2005. Biodiversity in model ecosystems I: coexistence conditions for competing species. – J. Theor. Biol. 235(4): 521–530.

Cheng, K.-S., Hsu, S.-B. and Lin, S.-S. 1982. Some results on global stability of a predator-prey system. – J. Math. Biol. 12(1): 115–126.

Del Giorgio, P. A. and Cole, J. J. 1998. Bacterial growth efficiency in natural aquatic systems. – Annu. Rev. Ecol. Syst. 29(1): 503–541.

Hassell, M. P., Lawton, J. H. and May, R. M. 1976. Patterns of Dynamical Behavior In Single-species Populations. – J. Anim. Ecol. 45(2): 471–486.

Hek, G. 2009. Geometric singular perturbation theory in biological practice. – J. Math. Biol. 60(3): 347–386.

Hofbauer, J. and Sigmund, K. 1998. Evolutionary Games and Population Dynamics. – Cambridge University Press.

Kuang, Y. 2002. Basic properties of mathematical population models. – J. Biomath. 17(2): 129–142.

Kuang, Y. and Freedman, H. 1988. Uniqueness of limit cycles in Gause-type models of predator-prey systems. – Math. Biosci. 88(1): 67–84.

López-Urrutia, A. and Morán, X. A. G. 2007. Resource limitation of bacterial production distorts the temperature dependence of oceanic carbon cycling. – Ecology 88(4): 817–822.

McPeek, M. 2012. Intraspecific density dependence and a guild of consumers coexisting on one resource. – Ecology 93(12): 2728–2735.

Pennekamp, F., Schtickzelle, N. and Petchey, O. L. 2015. BEMOVI, software for extracting behavior and morphology from videos, illustrated with analyses of microbes. – Ecol. Evol. 5(13): 2584–2595.

Rinaldi, S. and Muratori, S. 1992. Slow-fast limit cycles in predator-prey models. – Ecol. Model. 61(3-4): 287–308.

Schreiber, S. J. 1996. Global stability in consumer-resource cascades. – J. Math. Biol. 35(1): 37–48.

Schreiber, S. J. 2000. Criteria for C^*r*^ Robust Permanence. – J. Differ. Equ. 162(2): 400–426.

Thieme, H. R. 2003. Mathematics in Population Biology. – Princeton University Press.

